# Emergent representations of graphical structure in mechanistic neural models of causal judgment

**DOI:** 10.64898/2026.05.13.724819

**Authors:** Marcus A. Triplett, Kenneth Kay

## Abstract

Humans have a remarkable ability to judge causal relationships from a limited number of unreliable observations. Past work on causal cognition has largely focused on normative accounts of human behavior, leaving unknown how biologically plausible neural systems can learn causal relationships from observations and update representations of causal structure with additional evidence. Here, we leverage task-optimized recurrent neural networks to discover candidate implementation-level neural mechanisms of causal judgment. We propose a novel cognitive task in which a subject observes stochastic samples from an unknown causal structure (e.g. among variables *A, B*, and *C* with unknown causal relationships), and must judge whether a specific causal relationship is present given a query (e.g. “does *A* cause *C*?”). We found that, after training, recurrent neural networks perform the task at high accuracy, adopt strategies that incorporate observations from non-queried variables to form judgments, and exhibit neural activity with an emergent geometry reflecting underlying causal structure. We then use dynamical systems analysis to identify a set of low-level neural mechanisms that implement causal judgment and form representations of causal relationships. Together, these findings lay the groundwork for a “bottom-up” approach to causal cognition, providing a potential basis for experimental study in the brain.

## 1 Introduction

Humans routinely make judgments about causality based on observed events in the world. For example, recognizing whether a medication leads to symptom relief or whether a food type leads to stomach discomfort are intuitive judgments about cause and effect. These examples also highlight an important aspect of causal judgment in that they often require multiple observations: what if a headache resolved on its own rather than because of the medication? This process of reasoning about causation from a series of sparse and uncertain observations is ubiquitous in daily life [1], and has long been thought to be a cornerstone of human cognition [2]. Further, although we might often focus our attention on only a pair of variables (e.g. headaches vs medication), causal relationships are typically not judged in isolation. Humans commonly seek to understand the structure of causal relationships among several variables at once in order to consider possible alternative causes of an effect [3]. Importantly, this ability to consider causal interactions between multiple variables is thought to contribute to a range of cognitive functions, including imagining possible future or counterfactual outcomes [4, 3, 5–10], and guiding explanations [11, 12] and decisions [13–15].

Currently, the most widely used formalization of causal knowledge is the *directed acyclic graph* (DAG), in which directed edges between nodes represent causal relationships between objects or events in the world [16]. In particular, Bayesian networks (a probabilistic variant of DAGs) have become highly influential models in both psychology and machine learning. For example, there has been extensive research on estimating causal structure from observational data using Bayesian networks [17–20], and such approaches form the basis of leading cognitive theories of human causal induction [21, 4, 22–27] and causal inference in multisensory perception [28–32]. In Bayesian approaches to cognition, the brain is hypothesized to perform posterior inference over a limited number of possible causal structures, typically selecting the structure that best explains the observations, and potentially also updating its posteriors given additional evidence [33].

Although Bayesian approaches have been successful in accounting for a wide variety of human behaviors [34–37], it is unclear whether and how biologically plausible neural systems could implement the computations required by Bayesian causal model selection [38, 39, 30]. Further, it is currently not known how neural systems could in general infer causal relationships from experience, represent causal structure, and update such representations given additional experience. It is also worth noting that, due to implementation-level constraints, neural systems might adopt algorithms outside of those that have thus far been considered in cognitive science, which have instead focused largely on normative non-neural models of human behavior. Therefore, investigating the possible neural mechanisms of causal judgment could yield novel hypotheses at the algorithmic level of analysis [40], in addition to the implementation level.

Flexibly judging different causal relationships from a sequence of observations requires both integrating evidence over time [41–43] and performing context-dependent computations [44] (e.g. switching from judging whether *A* causes *B* to whether *B* causes *A*). Recurrent neural networks (RNNs) are among the simplest architectures with these abilities, and, crucially, neural mechanisms expressed in RNNs yield predictions that are testable in neural activity recorded from the brain [44, 45]. Thus, RNNs provide a natural model for investigating biologically plausible mechanisms of causal judgment. Here we study RNNs that have been trained on a novel causal judgment task in which the network observes stochastic samples from an unknown DAG and must judge whether a queried cause-effect relationship is present. We find that RNNs are capable of performing this task, incorporate observations of non-queried variables to resolve causal relationships, and exhibit neural activity that is geometrically organized to reflect the causal structure underlying the observations. Dynamical systems analysis [46–55] reveals putative neural mechanisms used to judge and represent causal relationships, together providing novel hypotheses that can guide future work probing the neural basis of causal cognition.

## 2 Training RNNs to make causal judgments

### 2.1 Preliminaries

Our approach is inspired by the structural causal models framework [16, 6, 56]. We specify a causal model as consisting of (1) a graph *G* with nodes {*A, B, C*, …*{* (representing distinct events or variables in the world) and directed edges {(*A, B*), (*C, B*), … *}*, together with (2) functions that assign each node a value (or *activity*) based on the values of the other nodes and any exogenous sources of variability. We say that a variable (represented by e.g. *X*) *causes* another (e.g. *Y* ) if there is a corresponding directed edge (*X, Y* ) in *G*. A node *Z* for which (*Z, X*) is a directed edge in *G* is referred to as a *parent* of *X*, and the set of all parent nodes of *X* is represented by pa(*X*). Finally, we focus on causal models whose directed edges do not form cycles (i.e., *DAGs*).

### 2.2 The DAG task

A canonical model for human causal judgment in cognitive science is the “noisy-or” parameterization of causal DAGs, where causes have independent chances of producing their effects [57, 4, 22, 58]. Formally, for a DAG *G* whose nodes *n* = *A, B, C*, … have activities *v*_*nk*_ at a sequence of observation times *k* = 1, …, *K*, samples from *G* are generated via ancestral sampling in the following manner. A node *n* can spontaneously activate with probability *p*_spont_ (the “leakage probability”), or can be activated otherwise by a parent node in *G* (if it has any) with probability *p*_cause_. Thus, the probability for node *n* to be active in observation *k* is a monotonically increasing function of the number of active parent nodes, given by

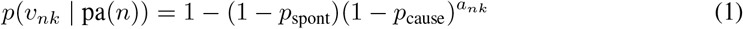

where *a*_*nk*_ = ∑_*m* pa(*n*)_ *v*_*mk*_ is the number of parent nodes of *n* active on observation *k*. Though in general DAGs are not uniquely identifiable without interventions [59, 56], the noisy-or parameterization introduces asymmetries in the observation statistics that improve identifiability of the underlying causal structure from observations alone while still being capable of expressing complex statistical dependencies [60]. For example, in a two-node DAG with one edge (*A, B*), the child node *B* is more active than the parent *A* – information that can be leveraged to resolve a causal relationship.

Here we propose to recast noisy-or structure learning as a *cognitive task*: a subject observes a sequence of activity from a collection of nodes with an unknown causal structure, together with a query regarding a specific causal relationship (“does *A* cause *B*?”), and must judge whether the query is true or false given the observations (Figure 1a). This task goes beyond simple binary evidence accumulation of instances of *A* and *B* because the subject can reason about the role of other variables (*C, D*, … ) that might confound a putative causal judgment. Further, the task goes beyond conventional context-dependent decision making [44, 61], as there are many more possible contexts (here causal queries).

**Figure 1:**
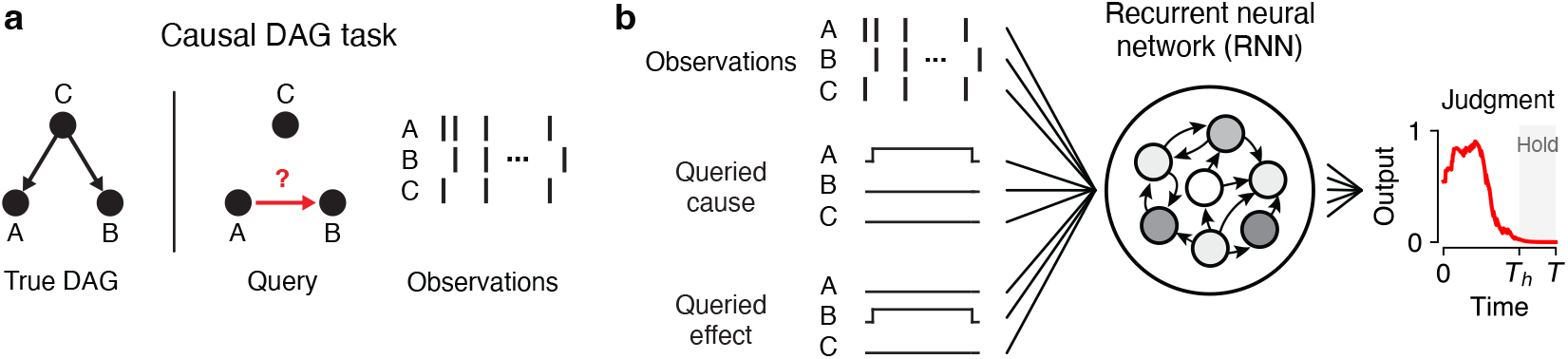
Causal judgment task and neural network model. **a**, Schematic of task. Observations are stochastically generated from a DAG whose true edges are unknown. The subject is queried (instructed or cued) on whether a causal relationship exists between a specific pair of nodes. **b**, Task implementation in neural networks. Inputs are a concatenation of three components: the node observations, the cause query (one-hot encoded), and the effect query (also one-hot encoded). RNNs are trained to report whether the queried causal relationship is true. Further details shown in Figure S1.

### 2.3 Recurrent neural networks

To discover putative mechanisms for how neural systems can form causal judgments, we used an approach in which RNNs were trained to solve the DAG task, then were reverse-engineered to identify their constituent dynamical computations. Importantly, this approach is agnostic regarding the process responsible for forming the causal judgment system itself (e.g. evolution, lifetime learning, or development [62]).

For the DAG task on *N* graph nodes, we considered nonlinear RNNs with hidden units that obey the following standard dynamical updates:

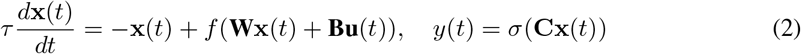

(e.g. see [63, 64, 52]; Figure 1b). Here **W** *∈* ℝ^*J×J*^ is the recurrent weight matrix, **u**(*t*) *∈* ℝ^*M*^ is the input vector at time *t*, **B** *∈* ℝ^*J×M*^ are the input weights, and *f* = tanh is a nonlinear activation function. Each **x**_*j*_ represents e.g. the firing rate of a neuron or putative collection of neurons. The RNN output at time *t* is represented by *y*(*t*), and computed from the hidden unit activity via output weights **C** *∈* ℝ^1*×J*^ followed by the logistic sigmoid function *σ*.

Inputs **u**(*t*) to the RNN consist of *M* -dimensional vectors made up of three concatenated segments (Figure 1b): the observed activity of the *N* graph nodes, the representation of the queried cause (one-hot encoded), and the representation of the queried effect (also one-hot encoded). Note that while node activity varies stochastically over time, the query inputs are tonically active. The RNN output corresponds to its judgment as to whether the queried cause-effect relationship exists in the underlying DAG. Importantly, the correct response to a given observation sequence is context-dependent, so the RNN must be able to reconfigure how it integrates the observations depending on the causal query. Further, it is worth emphasizing that, for a given observation sequence in the training data, the RNN only ever judges a single cause-effect query before those observations are discarded and the next observation sequence and query is generated (see Figure S1 for further details). Thus, RNNs are explicitly not trained to judge multiple different queries for any sequence of observations.

## 3 Results

### 3.1 RNNs can accurately perform the DAG task

We first focused on analyzing RNNs trained to make pairwise causal judgments on observations between three variables (*A, B, C*), as this is the smallest number of nodes in which non-trivial graph structures are possible: chains, forks, colliders, and mediators represent distinct causal relationships among three variables that do not appear in two-node systems (Figure 2a). Further, three-node DAGs are capable of expressing an extremely rich range of possible sequences of observations due to the diversity of graph structures and the stochastic nature in which nodes interact with each other.

**Figure 2:**
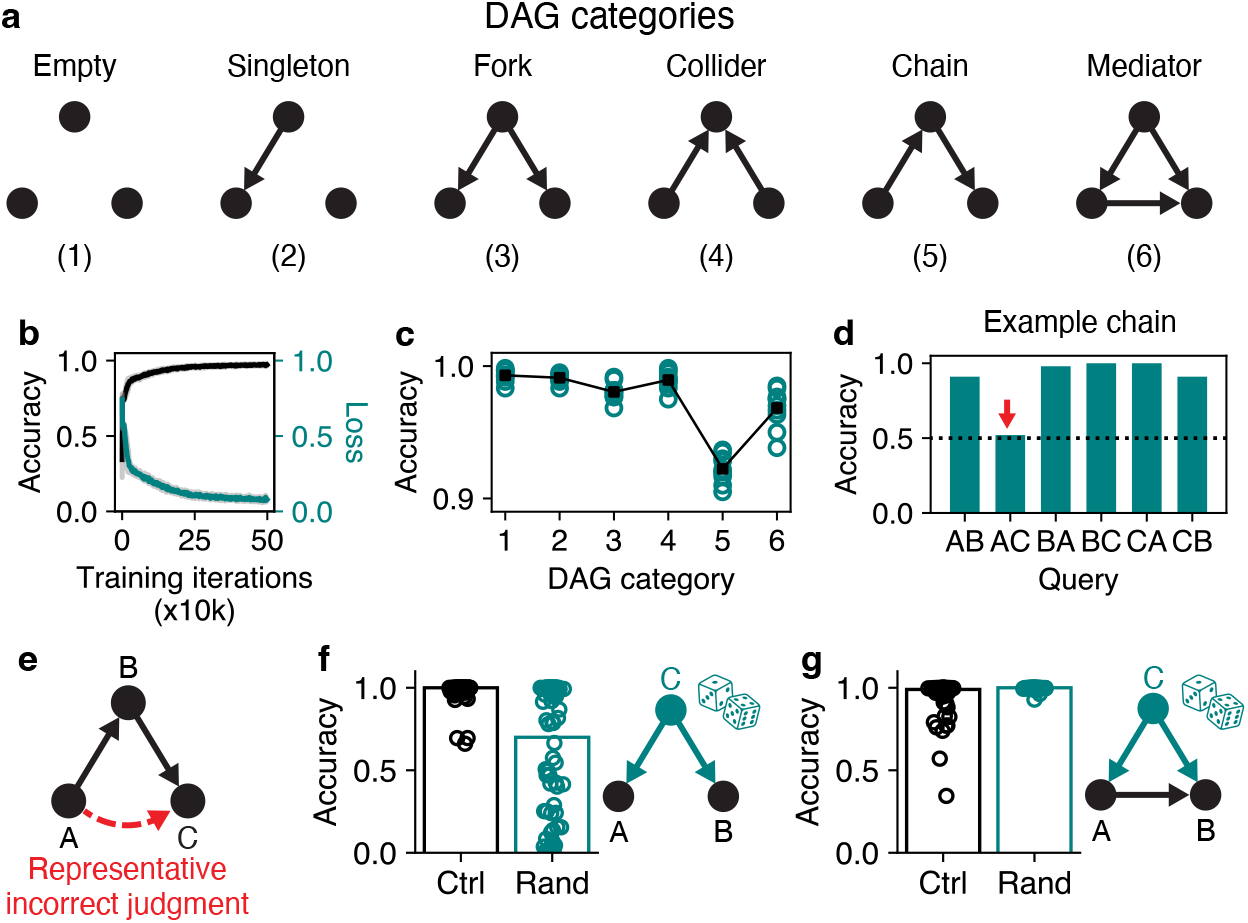
Trained RNNs can perform the DAG task and use non-queried variables to resolve causal ambiguities. **a**, The six categories of DAGs on three nodes. **b**, Accuracy and loss (on out-of-sample data) during training. Dark lines show averages over 10 randomly initialized RNNs. Faint gray regions show mean*±* 1 s.d. **c**, Accuracy of trained RNNs as a function of DAG category. Each data point represents the average performance of an RNN on 100 randomly generated DAGs of the listed category. **d**, Accuracy of an example trained RNN as a function of all possible queries on a chain DAG. While the RNN achieves high performance on five out of six queries, query *AC* (“does *A* cause *C*?”, red arrow) is at chance level (dotted line). **e**, The reduced accuracy on chain DAGs arises from the RNN often inferring that *A* causes *C* directly when in fact *A* causes *B* and then *B* causes *C* (and similarly for other permutations of *A, B*, and *C* in chain DAGs). **f**, Accuracy of the RNN on judging whether *A* causes *B* for a fork DAG (right), under normal conditions (“Ctrl”, black circles) vs. when the activity of *C* is randomized (teal circles). Randomizing *C* significantly lowers RNN accuracy (*p <* 10^*−*17^, Wilcoxon signed-rank test), often causing it to report that *A* does cause *B* since the covariance of *A* and *B* can no longer be explained by another variable. **g**, Similar to (f), but where *A* does cause *B*. Randomizing *C* improves RNN accuracy (*p <* 10^*−*15^).

After training, we found that RNNs performed the task with high overall performance, achieving a mean accuracy of 0.95 (out of 1) on novel observations sampled from three-node DAGs (Figure 2b).

This indicates that RNNs can successfully judge cause and effect in the DAG task, and thus provides a starting point for analyzing mechanisms of causal judgment.

### 3.2 Strategies and failure modes in trained RNNs

Despite achieving high accuracy, RNNs still made errors, prompting us to investigate possible behavioral biases. To do so, we categorized all 25 DAGs on 3 nodes into 6 distinct categories: the empty DAG, DAGs with a single edge, fork DAGs, collider DAGs, chain DAGs, and mediator DAGs (Figure 2a). Of these categories, RNNs had markedly lower accuracy on chain DAGs in particular (Figure 2c), and, more specifically, tended to erroneously judge that *A* directly causes *C* in circumstances where in fact *A* causes *B* and *B* then causes *C* (Figure 2d,e). Thus, trained RNNs displayed a transitivity bias, similar to human causal judgment under some conditions [65–67].

Next we sought to probe the underlying strategy adopted by the RNNs. If an RNN is judging whether *A* causes *B*, to what extent does it care about *C*? To evaluate this capacity, we considered a fork DAG in which the only causal effects are that *C* causes *A* and *C* causes *B*, and then queried the RNN as to how it judges if *A* causes *B*. This case is potentially insightful because the RNN might ignore *C* and selectively integrate instances of *A* without *B*. Alternatively, the RNN could leverage the activity of *C* to correctly identify it as a common cause of both *A* and *B*, despite not being explicitly queried on the role of *C*.

To resolve these alternatives, we randomized the activity of node *C* (via circular shuffles in time) while leaving the activity of *A* and *B* intact. Intriguingly, doing so led to a substantial decrease in RNN performance (Figure 2f), indicating a critical role for observations of *C* when judging whether *A* causes *B*. Further, we also simulated observations from a DAG in which *A* acted as a mediator between *C* and *B* (i.e. *C* causes *A, C* causes *B*, and *A* causes *B*). We found that the RNN’s ability to discriminate the causal effect of *A* on *B* had a small but statistically significant improvement when *C* was randomized (Figure 2g), presumably due to *C* presenting as less of a potential confounder under randomization. Together, these results show emergent utilization of information about non-queried variables to judge causal relationships, as observed in human causal reasoning [57].

We next performed a more stringent test: whether RNNs can accurately judge the entire graphical structure rather than an individual edge in isolation. To that end, we sampled a single sequence of observations from an example DAG, and then cycled through every possible causal query with the observations kept fixed. We found that, despite only ever being trained on a single cause-effect query per observation sequence, RNNs can indeed resolve entire graphs (Figure 3a-c, Figure S2).

**Figure 3:**
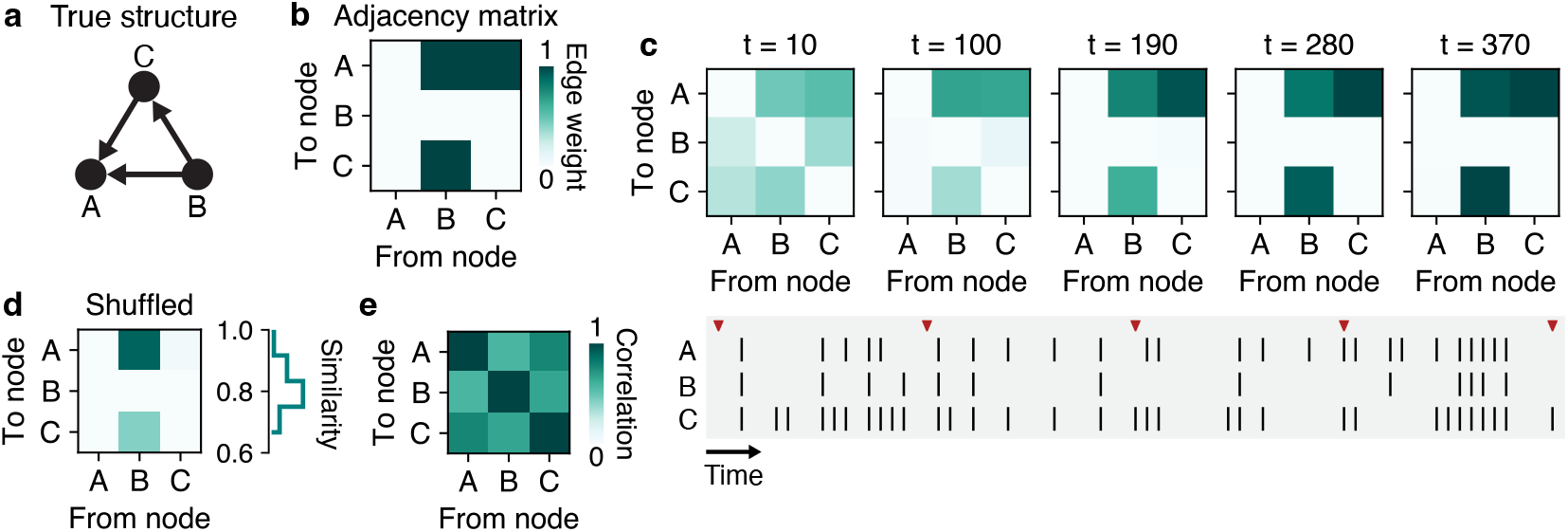
Testing RNNs on the complete graphical structure underlying observations. **a**, Example DAG on which behavior was tested across queries for a single fixed observation sequence. **b**, Adjacency matrix corresponding to the DAG in (a). **c**, Top: Evolution of an RNN’s judgments on all causal relationships over the course of a trial. Bottom: Sequence of observations used to form the judgments made in top panel. Red ticks correspond to the times sampled. **d**, Control analysis on marginal statistics. Left: shuffling (via cyclic permutations) the marginal activity of each node alters the inferred graphical structure, here eliminating *C* to *A* (single example shown). Right: similarity between the graphical structure inferred from the actual vs. shuffled observations (distribution over 100 random shuffles shown). Similarity defined as the fraction of graph edges inferred from the shuffled observations that are equal to the graph edges inferred from the actual observations. **e**, Correlation coefficients (derived from observations in (c)) between every pair of nodes shows that RNNs do not simply report correlations. Additional examples shown in Figure S2.

Lastly, we asked whether the RNN was simply integrating the marginal statistics of each node’s activity to form its judgments. As noted in subsection 2.2, the noisy-or parameterization can induce signatures of causal relationships in the marginal statistics of each node. For example, a node with no parent nodes can only activate via the leakage probability – a relatively rare occurrence compared to if the node had several parent nodes, each capable of activating the child node. We conducted a shuffle analysis by performing a circular permutation independently for each node’s observation sequence, such that the covariance between node activity was eliminated, but the marginal activity statistics were all preserved. Under this analysis, we found that RNNs failed to recover the true DAG structure, indicating that they were not simply making judgments via the marginal statistics (Figure 3d)^1^. Similarly, we found that RNNs were not simply reporting the correlation structure of the observations (Figure 3e), which could potentially show similarity to the causal structure. Thus, trained RNNs formed their judgments via both marginal and covariance statistics.

### 3.3 RNNs generalize to unseen classes of graphs

We next tested whether trained RNNs had the capacity to generalize their judgments to entire classes of graphs not seen in training. We focused on the DAG task with 4 nodes, which yields a large and diverse collection of graph structures while retaining computational tractability. On 4 nodes there are 543 DAGs, which can be partitioned into 31 distinct classes (up to isomorphism). We trained RNNs on 25 graph classes, tested on the 6 remaining (out-of-distribution) classes, and then cycled through which classes were designated as training versus testing. We found that RNNs performed with an accuracy of 0.93 *±*0.04 on novel graph classes (*±*1 s.d., *n* = 5 randomly initialized RNNs tested per cross-validation fold), confirming that RNNs can indeed generalize to unseen causal structures (Figure S4, Figure S5).

### 3.4 RNN activity shows an emergent geometry reflecting graphical structure

Towards understanding how neural systems actually perform causal judgments, we next examined the activity trajectories of a representative RNN. For visualization, we projected the trajectories of the hidden units onto the top three principal components (PCs), which together accounted for 75% of the variance for this RNN (Figure S6). There are six possible causal queries for a three-node DAG (*A* causes *B, A* causes *C, B* causes *A*, etc.). As such, we first simultaneously plotted the trajectories for each of the six possible queries in the absence of any observations. These plots showed distinctive structure: trajectories radially diverged from a common starting point near the origin before settling at query-dependent locations that were approximately equidistant from their neighbors (Figure 4a). Hidden states showed minimal movement thereafter, suggesting that recurrent dynamics guided trajectories towards stable fixed points associated with each query.

**Figure 4:**
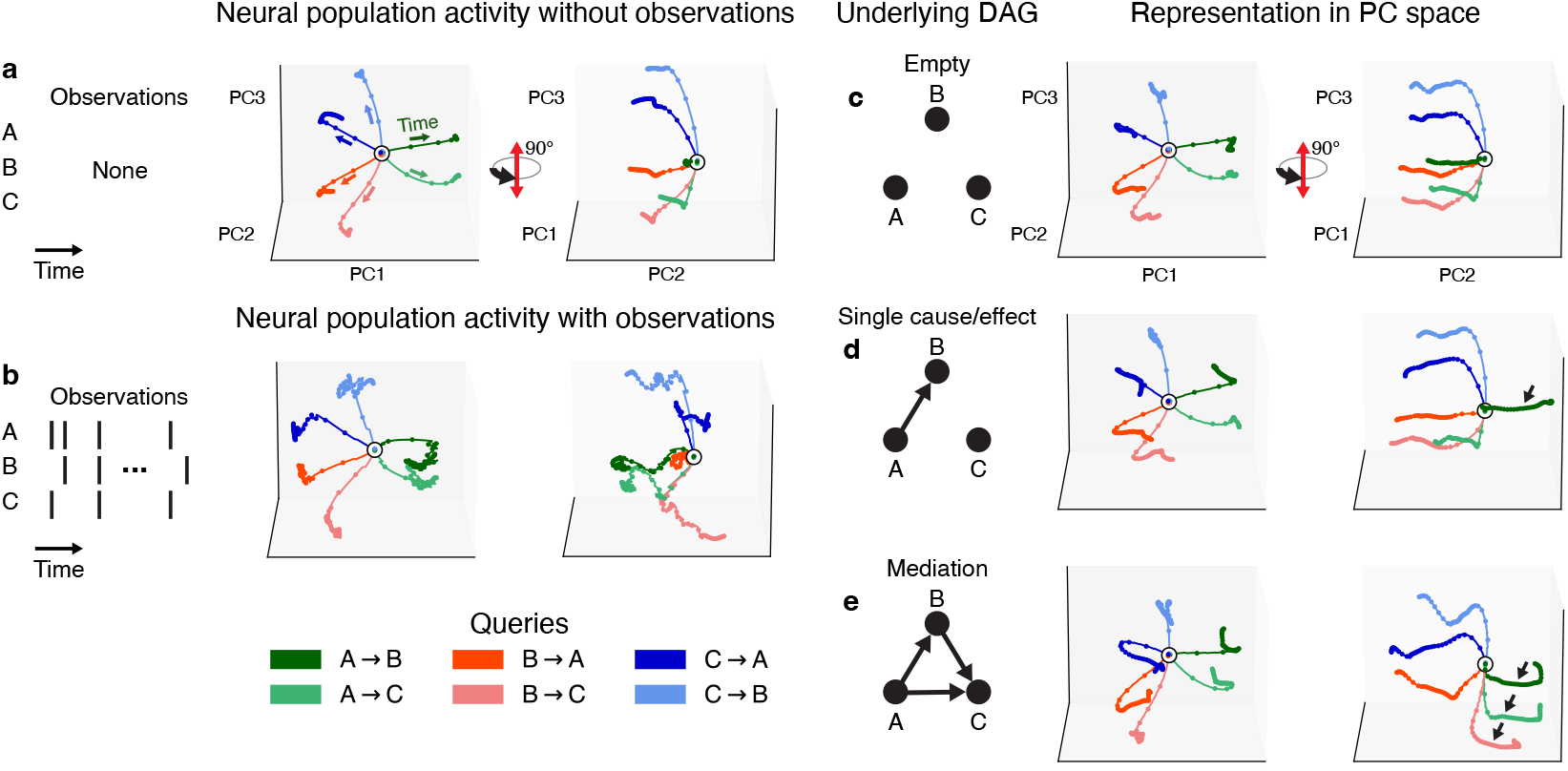
Emergent representations of graphical structure in trained RNNs. **a**, Neural trajectories generated by a trained RNN in the absence of observations (but with tonic query input enabled). Colored paths correspond to different queries. Right panel rotated 90^*°*^. **b**, Same as (a),but with observations sampled from an example DAG. **c-e**, Average neural trajectories correspond-ing to DAGs with increasing numbers of edges. Averages taken over 500 samples of observation sequences. Black arrows in plots in (d) and (e) indicate trajectories corresponding to queries that are true in corresponding DAG. All panels and trajectories plotted in common PC space.

Next, we sampled a single sequence of observations from an example DAG (*B* causes *C, C* causes *A*), constituting a single trial of the DAG task, and performed a similar visualization. The corresponding trajectories were similar to the observation-free case in that they diverged from a common initial point, but were considerably more irregular due to the stream of stochastic inputs (Figure 4b).

We then began investigating the representation of specific causal graphs. We started by sampling observations from the empty DAG (such that nodes only activate spontaneously) and visualizing the average trajectory over 500 trials. (Note that by a *trial* we mean a sequence of observations from a DAG, along with a query input). This recapitulated the radial divergence seen in the observation-free case, but with elongated trajectories associated with “false” RNN outputs (Figure 4c). However, as we introduced edges into the underlying DAG and further examined activity across trials, we found that the associated trajectories traced paths in the opposite direction in state space compared to when the edge did not exist (Figure 4d,e). Averaging trajectories with causal relationships being present (edge exists) or absent (edge does not exist) yielded an effective decision axis that ran parallel to the RNN trajectories and orthogonal to a decision plane (Figure S7).

These findings implied a systematic implementation for representing arbitrary causal graphs, where query-specific trajectories move towards either side of a decision plane depending on how the RNN integrates the input sequence.

### 3.5 RNNs use query-specific line attractors to represent causal relationships

To go beyond visualization of the hidden unit trajectories, we next sought to uncover the specific dynamical mechanisms underlying the formation of causal judgments. To do so, we adopted a “reverse engineering” approach to understanding RNNs [46, 44, 47]. Briefly, the approach consists of identifying fixed points in the RNN dynamics (i.e. neural states where *d***x**(*t*)*/dt ≈*0), characterizing how the fixed points relate to RNN output, and understanding how the neural state is driven between fixed points via interactions between the RNN’s external inputs and its recurrent dynamics. In particular, we searched for fixed points while observations were absent but the query input was on, thereby simulating observation-free periods in the task (see Appendix B for details).

We found that fixed points were organized into query-specific line attractors, all of which were aligned with the decision axis (Figure 5a,b). Further, this fixed point structure was also the case for RNNs trained to judge causal effects in DAGs with two, three, or four nodes (Figure S8). Linearization of the RNN dynamics around each fixed point revealed that the corresponding largest real-valued eigenvalues were approximately equal to 1 (Figure S9), consistent with a stable integrating mode (Figure S10). Moreover, line attractors showed a striking organization – they were clustered in space according to the queried cause: the *A →X* line attractors were closer together and had visibly more similar dynamics compared to the *B→ Y* and *C→ Z* line attractors (Figure 6a-c), indicating a systematic organization of population activity relative to the causal graph.

**Figure 5:**
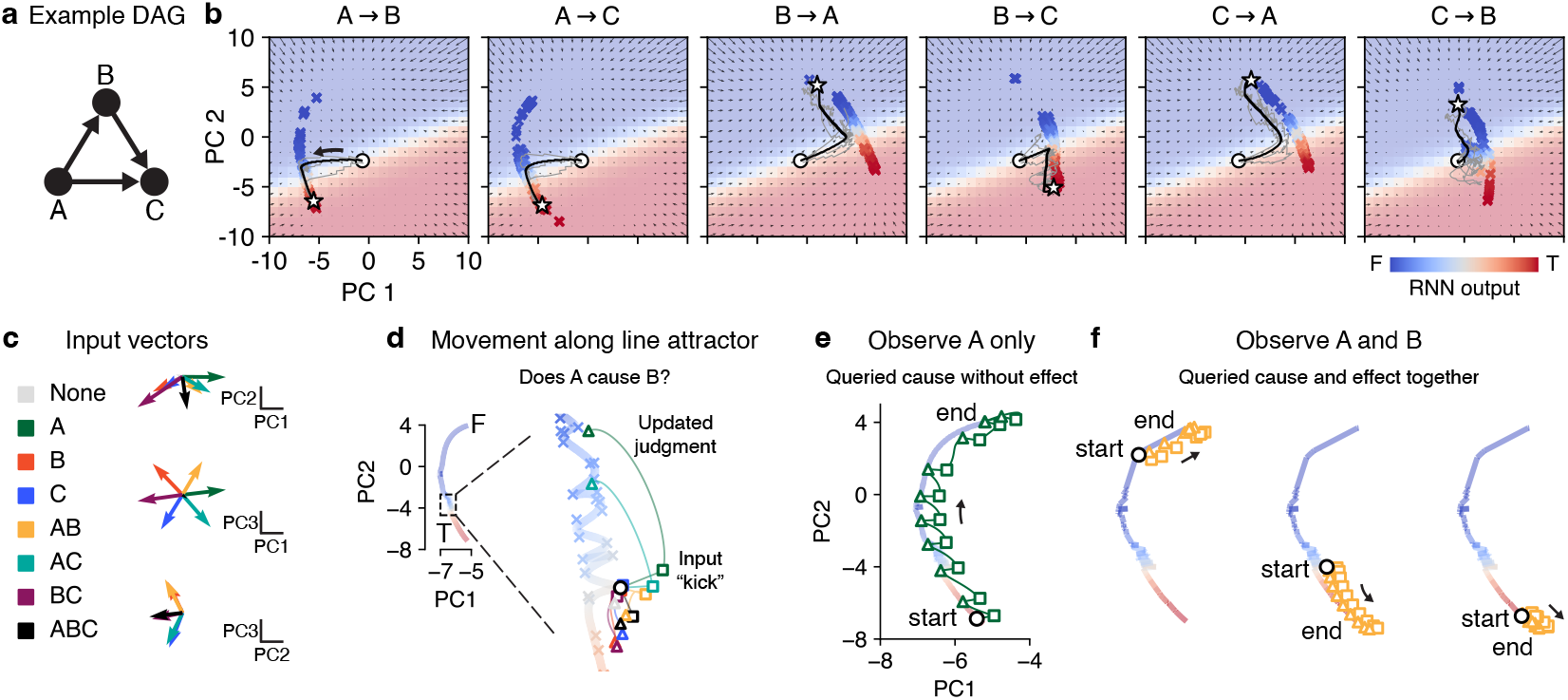
Formation of causal judgments via line attractor dynamics. **a**, Example DAG used to generate observations. **b**, Fixed points (colored crosses) are organized into an approximate line attractor for each query (specified above each panel). Vector field shows approximate recurrent dynamics (in top two PCs) for each query. White circles: initial states; white stars: final states; black lines: average trajectories over 500 trials sampled from DAG in (a); faint gray lines: example trajectories from three trials. See Figure S8 for examples for other node counts. **c**, Left: color legend for all eight possible observations (*A* alone, *B* alone, *AB, BC*, etc). Right: input vectors 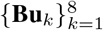 corresponding to all possible observations, projected onto top three PCs. **d**, Interaction between input vectors and recurrent dynamics differentially moves neural state along the line attractor (represented by thick colored line interpolating between fixed points). Colored squares show instantaneous effects of different inputs. Colored triangles show neural states after 200 timesteps. **e**, Repeated observations of *A* induce the neural state to traverse line attractor from end to end. **f**, Similar to (e), but for observing *A* and *B* together. Note the dependence of outcome on initial state.

**Figure 6:**
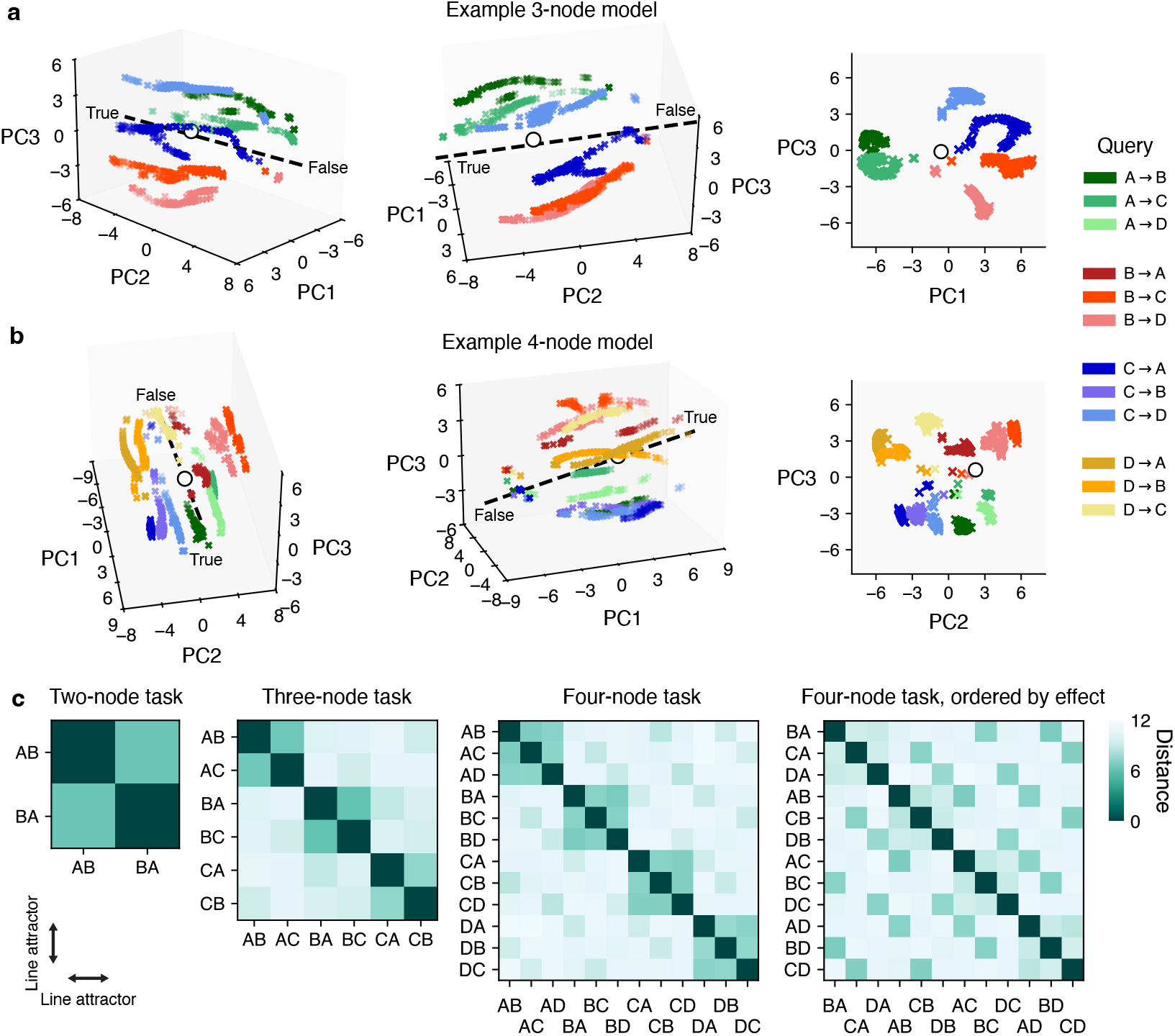
Clustering of line attractors (“judgment axes”) across queries. **a**, Visualization of all line attractors simultaneously for an RNN trained on the 3-node DAG task. Line attractors colored by query. Crosses represent fixed points; dashed line, output axis; white circle, coordinates of initial state. **b**, Same as (a), but for an example RNN trained on the 4-node DAG task. **c**, Euclidean distances between line attractor centroids across RNNs trained on the DAG task with 2, 3, and 4 nodes. Each data point represents an average over 10 randomly initialized RNNs. First three panels ordered to highlight that clustering occurs primarily via the queried cause. Rightmost panel re-ordered to show secondary clustering occurs with respect to the queried effect.

### 3.6 Probing input-specific dynamics of updates to causal judgments

We next sought to understand how different observations moved the neural state along the line attractors. As an example, we focused on the *A→ B* query, though the core mechanisms hold more generally. We selected a fixed point in the middle of the line attractor and visualized the effect of every possible input (*A* alone, *A* and *B* together, *B* and *C* together, etc). Different input combinations displaced (“kicked”) the neural state in different directions according to the instantaneous effect of the input vectors (Figure 5c,d). Then, depending on the input-driven repositioning, recurrent dynamics carried the neural state back to the line attractor, but further towards the true or false ends depending on whether the observation supported or contradicted the query (Figure 5d).

This “kick-and-roll” activity pattern suggests a mechanism through which the RNN can traverse the line attractor in order to make its judgment. For example, when querying whether *A* causes *B*, repeatedly observing *A* alone iteratively kicks the neural state towards the “false” end of the line attractor (it is unlikely that *A* causes *B* if one often observes *A* without also *B*), regardless of where the neural state starts (Figure 5e). On the other hand, observing *A* and *B* together supports the query, and therefore typically moves the neural state towards the “true” end of the line attractor (Figure 5f, middle and right). Interestingly however, this effect is state-dependent along the line attractor: observing *AB* supports *A* causing *B* unless the RNN already “believes” *A* does not cause *B*, in which case all observations (including *AB*) move the neural state even further towards the “false” end (Figure 5f, left). Finally, we comprehensively quantified the extent to which every possible observation moves the neural state along each query’s line attractor starting from every fixed point, revealing a diversity of state-dependent input effects that together describe how the RNN makes its judgments (Figure S11).

#### 3.7 A neural algorithm for forming causal judgments

Given these results, we derived a mechanistic model of how recurrent neural circuits can judge and represent causal relational structure (Figure 7). Our model is based on the linearized RNN dynamics: first, using a first-order Taylor series, the local dynamics around a fixed point **x**^*\**^ on the line attractor can be approximated as a linear dynamical system **z**_*t*+1_ *≈* **J**_rec_**z**_*t*_ + **J**_inp_**w**_*t*+1_, where **J**_rec_ and **J**_inp_ are the Jacobians of the RNN update function with respect to the neural (hidden) state and the input, and where **w**_*t*_ is an input term (see Appendix C for details). The long-term behavior of the neural state (e.g. for *t* = 1, …, *T* ) in response to a single observation **ŵ**can be approximated in the neighborhood of **x**^*\**^ in terms of the slowest eigenmode as **z**_*T*_ *≈* **r**(***l***^*⊤*^**J**_inp_**ŵ** ), where **r** and ***l***^*⊤*^ are the right and left eigenvectors associated with the largest eigenvalue of **J**_rec_ (in our RNNs this is typically *λ*_1_ *≈* 1). The expression ***l***^*⊤*^**J**_inp_**ŵ** is a scalar that describes how far the neural state will move in direction **r**, as a function of how the input vector **J**_inp_**ŵ** aligns with left eigenvector ***l***^*⊤*^.

**Figure 7:**
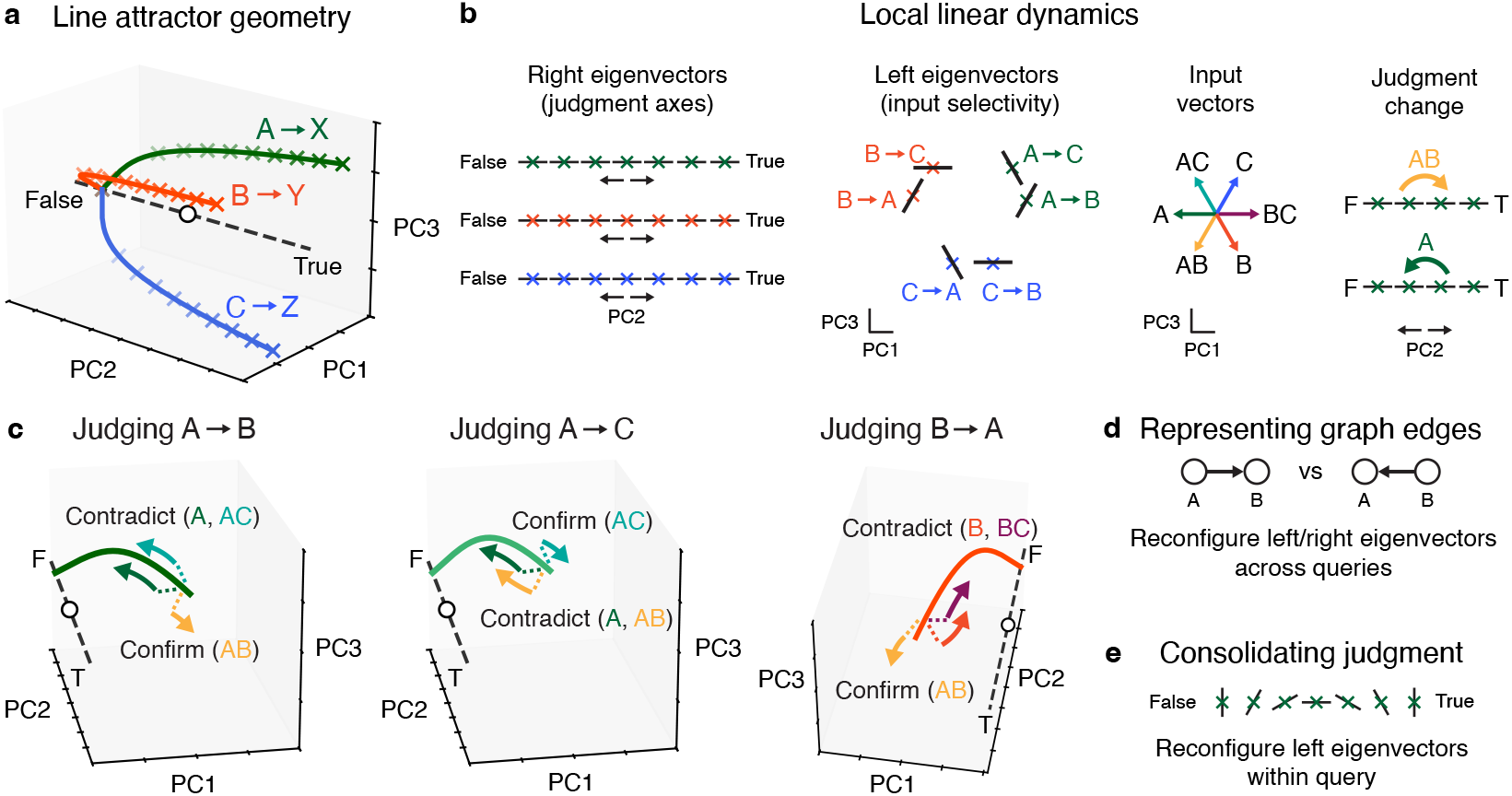
Population dynamics and geometry for a neural implementation of causal judgment. **a**, Stylized line attractors for each causal query, clustered by the queried cause. Colored crosses represent fixed points; dashed line, trained output weights plotted as direction in PC space; white circle, neural activity state at trial start. **b**, Local dynamics underlying how neural state transitions between fixed points on a line attractor. For each query, associated fixed points and their right eigenvectors define a stable “judgment axis” ranging from “false” to “true”. Left eigenvectors determine input selectivity: an input aligned with a left eigenvector will drive the neural state towards “true”, whereas an anti-aligned input will drive state towards “false”. **c**, Three examples of how local recurrent dynamics are reconfigured to enable opposing effects of inputs under different queries (see also Figure S12). **d**, Representation of graph edges implemented via reconfiguration of left and right eigenvectors *across* queries. **e**, Left eigenvectors also reconfigure *within* queries, becoming orthogonal to the inputs, possibly as a mechanism to consolidate judgments.

For a given query, a line attractor is composed of a collection of such fixed points (Figure 7a), each of which is associated with a right eigenvector **r** specifying a direction of movement in state space. Collectively, these points define a “judgment axis” that spans from “false” to “true” (Figure 7b). By orienting the corresponding left eigenvectors such that they align *specifically* with observations of the cause and effect together, but are anti-aligned with observations of the cause without the effect, the neural state will move appropriately towards the true or false ends of the judgment axis.

However, when the queried graph edge changes (e.g. from *A →B* to *A→ C*), the recurrent dynamics must reconfigure accordingly: an observation of *AC* switches from being contradictory evidence to confirmatory evidence (Figure 7c). Thus, *using the same recurrent connectivity matrix*, a change in query induces a reconfiguration of the (locally linear) recurrent dynamics such that the left eigenvector switches from being anti-aligned with observation *AC* to being aligned with it (Figure 7d).

Further, clustering line attractors by the queried cause (Figure 6c) allows the RNN to share local dynamics across queries: in particular, observing the cause without the effect drives the neural state towards the false end of the corresponding line attractor, regardless of which specific effect is being queried. Finally, the state-dependent effect of an observation *within* queries (Figure 5f) is consistent with an additional reconfiguration of the left eigenvectors within the line attractors, where, at the extreme ends of each line attractor, the left eigenvectors become orthogonal to (or even anti-aligned with) observations they were previously aligned with (Figure 7e). This reconfiguration could constitute a mechanism to consolidate a judgment.

## 4 Conclusion

Here we identified candidate neural mechanisms of causal reasoning by reverse-engineering RNNs trained on a novel cognitive task. Our results show that, rather than explicitly performing model selection over causal graphs [22], neural systems could judge and represent causal structure via emergent neural dynamics and geometry. This provides a mechanistic alternative to conventional normative accounts of human causal judgment [38]. Our results also connect to neural geometry in multitask RNNs [68–70], since each causal query in our task is effectively its own task. Importantly, the number of possible queries grows quadratically with the number of graph nodes, requiring numerous query-specific reconfigurations of how evidence is integrated, and thus favors separation between line attractors in state space to limit interference. At the same time, symmetries in the DAG task enable particular dynamical motifs to be shared across queries [52] (e.g. observing *A* alone should affect judgment similarly across queries *A →B* and *A→ C*), favoring line attractors being clustered by their queried cause. These competing pressures appear to yield a hierarchical geometry where line attractors are organized primarily by cause and secondarily by effect (Figure 6), in turn enabling compositional reuse of dynamical motifs across causal queries.

## Acknowledgements

The authors would like to thank Larry Abbott, John Pearson, Benjamin Antin, and Samuel Lippl for helpful discussions and feedback. MAT is supported by NIH award K99NS135649, the Gatsby Charitable Foundation (award GAT3708), and the Kavli Foundation.

## Appendix

### A Recurrent neural network training details

#### A.1 Network and optimization parameters

We trained RNNs with *J* = 128 hidden units with tanh nonlinearities using the cross-entropy loss function (Figure S1). We used a time constant corresponding to *τ* = 13 time steps. RNNs were trained to hold their output for a period of 179 time steps. An RNN output with *y*(*t*) *≥* 0.5 represents it making a “true” judgment, and otherwise a “false” judgment. Both the recurrent weights and the hidden unit activity were regularized using L2 penalties with coefficients *λ*_rec_ = 10^*−*4^ and *λ*_hid_ = 10^*−*5^. RNNs were trained with a batch size of 1024. During training, hidden unit activations were injected with Gaussian noise (mean, 0; standard deviation, 0.025) to improve out-of-sample performance. Sets of 10 RNNs were trained for 25,000, 50,000, or 75,000 epochs for DAG tasks with 2, 3, or 4 nodes respectively, which was sufficiently many epochs to obtain RNNs that perform the task with high accuracy and interpretable dynamical structures. RNNs were trained using the Adam optimizer with a learning rate of 4 *×* 10^*−*4^. We also found that RNNs could be successfully trained on DAGs with higher node counts (accuracy: 0.92, 0.87, and 0.85 for the DAG task with 5, 6, and 7 nodes respectively; means taken over 5 randomly initialized RNNs; number of training iterations fixed at 100,000).

#### A.2 Generating training data

In our training data, a sequence of observations was made up of 75 samples from a DAG, separated in time by an interstimulus interval of 5 time steps. Nodes in the DAG probabilistically activated each other under the noisy-or rule described in subsection 2.2, where *p*_cause_ = 0.8 and *p*_spont_ = 0.2. To construct a DAG, we first initialized its adjacency matrix with all zeroes, and then (to ensure acyclicity) filled in the upper triangle only, sampling binary values uniformly at random. Node labels were then randomly permuted. Consequently, for a fixed pair of nodes (*A, B*), the marginal probability that query *A→ B* is true across all randomly sampled DAGs in the training data is 0.25. During training, networks are presented with all possible *N* -node DAGs, but only a small fraction of all possible sequences of observations. For example, although there are only 25 possible DAGs on three nodes, it is possible to generate extremely large quantities of training and test data by stochastically sampling different observation sequences. In our case, since we trained our RNNs using observation sequences comprised of 75 temporally-spaced samples, there are 2^225^ possible sequences for each DAG. We also verified that RNNs generalize to held-out (“unseen”) classes of DAGs (see subsection 3.3).

### B Fixed point analysis

To identify fixed points, we minimized a discrete-time analogue of the “*q* function” from ref. [46]:

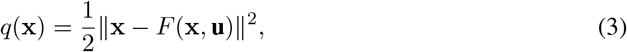

where *F* represents the one-step update resulting from discretizing the continuous RNN dynamics in Equation 2 (see Equation 6 below). Here we have suppressed the dependence on time to simplify notation. We focused our analysis on minimizers **x**^*\**^ of Equation 3 such that *q*(**x**^*\**^) *<* 10^*−*5^, and thus we typically identified approximate fixed points, or “slow points” [71].

Based on the behavior of the RNN when simulated without observations (subsection 3.4), we hypothesized that any fixed point structure would be query-specific (as the query input is tonically on in a trial). When searching for fixed points, we therefore provided a static input vector **ũ**_*k*_ *∈* ℝ^*M*^ for which the node activity is zero but for which query *k* is active; i.e. *ũ*_*ki*_ = 0 for *i* = 1, …, *N*, and *ũ*_*ki*_ = *ũ*_*kj*_ = 1 if *i* and *j* correspond to the queried cause and effect indices respectively, and *ũ*_*ki*_ = 0 otherwise.

To seed the fixed point finding algorithm with initial activity states, we used the following procedure. When the number of nodes was three or less, we enumerated all possible DAGs, simulated RNN trajectories on each DAG, and sampled initial states at three time steps along each average trajectory, as well as from two randomly selected single trials. When the number of nodes was larger than three, the number of possible DAGs grew extremely quickly, making the number of possible seeds too large to test. Instead, we sampled initial states from the empty DAG, a complete DAG (where there is an edge of some orientation between every pair of nodes, e.g. {(*A, B*), (*A, C*), (*A, D*), (*B, C*), (*B, D*), (*C, D*) }), the complete DAG with all edges reversed, as well as the singleton DAGs (i.e. the DAG with exactly one causal edge, as in Figure 2a, for every possible edge). This ensured that the fixed point finder was seeded with a variety of “true” and “false” judgments for every possible causal query.

### C Linear dynamical systems analysis

In this section we derive the linear approximation to the query-specific RNN dynamics, much of which follows the derivations from refs. [44, 50]. First, recall that the continuous-time RNN dynamics are given by

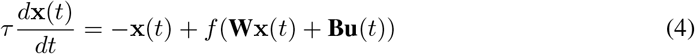

where *f* is the tanh activation function, and where the RNN output is subsequently obtained as *y*(*t*) = *σ*(**Cx**(*t*)). In practice, we run the RNN by approximating the continuous-time dynamics via the following discrete-time update rule:

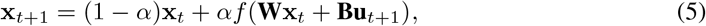

where *α* = 1*/τ* . (Note that Equation 5 is obtained from Equation 4 via the finite difference approximation).

Next, we wish to understand the dynamics of the discrete-time RNN around a fixed point **x**^*\**^ associated with query *q* (where *q* indexes the causal queries *A →B, A→ C*, etc.). To that end, we define the RNN update function *F* as

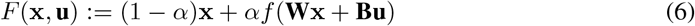

and set Δ**x**_*t*_ = **x**_*t*_ *−* **x**^*\**^ (i.e. the neural state relative to the chosen fixed point) and Δ**u**_*t*_ = **u**_*t*_ *−* **u**^*\**^. Since we are interested in query-specific dynamics, we choose **u**^*\**^ *∈* ℝ^*M*^ to be the input correspond-ing to query *k* but without observations, analogous to the input used during fixed point finding. Then (following e.g. ref. [50]) we perform a first-order Taylor approximation about the pair (**x**^*\**^, **u**^*\**^):

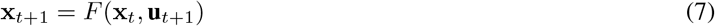

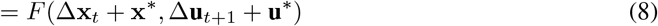

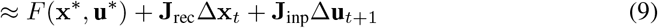

where [**J**_rec_]_*ij*_ = *∂F* (**x**^*\**^, **u**^*\**^)_*i*_*/∂***x**_*j*_ and [**J**_inp_]_*ij*_ = *∂F* (**x**^*\**^, **u**^*\**^)_*i*_*/∂***u**_*j*_ are the Jacobians of the dynamical update function with respect to the neural state and the input respectively. Next, because **x**^*\**^ is a fixed point of *F* associated with query **u**^*\**^, *F* (**x**^*\**^, **u**^*\**^) *≈* **x**^*\**^. Hence we obtain the following local linear approximation to the RNN dynamics:

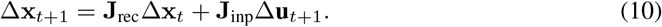

Finally, setting **z**_*t*_ := Δ**x**_*t*_ and **w**_*t*_ := Δ**u**_*t*_, we have the linear dynamical system from subsection 3.7,

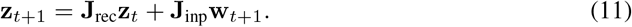

A powerful approach to evaluating the long term behavior of a linear dynamical system is in terms of its eigendecomposition. In our linearized RNNs, the recurrent dynamics matrices are generally *non-normal* [72–74]; i.e., 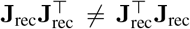. Importantly, for non-normal matrices the eigende-composition is defined in terms of both left and right eigenvectors,

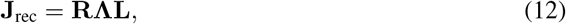

where **Λ** *∈* ℝ^*J×J*^ is a diagonal matrix whose diagonal terms are the eigenvalues *λ*_1_, …, *λ*_*J*_ (ordered from largest magnitude to smallest), **R** *∈* ℝ^*J×J*^ is a matrix whose columns **r**_1_, …, **r**_*J*_ are the right eigenvectors of **J**_rec_, and **L** = **R**^*−*1^ is a matrix such that each row 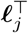 is a left eigenvector associated with the right eigenvector **r**_*j*_. To evaluate the long term behavior of **x** (e.g. for some large *T* ) in response to a single observation **û** near fixed point **x**^*\**^, we consider the linearized system with **ŵ** = **û** *−* **u**^*\**^,

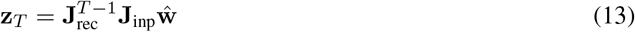

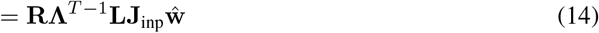

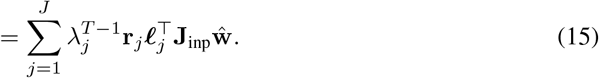

In our RNNs, the leading eigenmodes are either associated with a single real-valued eigenvalue near 1, or come in complex conjugate pairs (Figure S9). In the latter case, these modes generate partial oscillatory activity patterns within two-dimensional planes to induce movement along the line attractor. Here, we restrict our focus to the former case (i.e. the real eigenmodes) as these provide a simple and sufficient implementation-level description of how a linear recurrent neural system can judge causation within the context of a specific query. Under this assumption, the eigenmodes with eigenvalues *<* 1 decay rapidly and thus, for sufficiently large *T*, the long term behavior of **z** is approximated by the expression

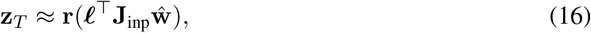

where we have dropped the subscripts on the left and right eigenvectors since we assume that they correspond to the eigenvector with largest eigenvalue. Further, we suppress reference to the eigenvalue itself since it is assumed to be approximately equal to 1, and therefore 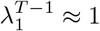.

The term ***l***^*⊤*^**J**_inp_**ŵ** is a scalar that describes how far **z** will move in direction **r** as a function of the alignment between the input vector **J**_inp_**ŵ** and the left eigenvector ***l***^*⊤*^. This relationship between the inputs and the left and right eigenvectors plays a crucial role in our proposed neural implementation of causal judgment. If the input vector is aligned with the left eigenvector, then ***l***^*⊤*^**J**_inp_**ŵ** *>* 0, and **z** will move in direction **r** (i.e. towards the “true” end of the judgment axis)^2^. If the input is anti-aligned with the left eigenvector, (i.e. ***l***^*⊤*^**J**_inp_**ŵ** *<* 0) then **z** will move in direction *−***r** (towards “false”). Additionally, the RNN can selectively ignore an input by having the left eigenvector oriented such that ***l***^*⊤*^**J**_inp_**ŵ** = 0 (i.e., oriented orthogonally). See Figure 7 and Figure S12 for visual schematics providing further intuition about the interaction between left eigenvectors and inputs.

Because an observation can switch from being confirmatory to contradictory when the query is changed, the left eigenvectors must be reconfigured (i.e. reoriented) by the query input to facilitate the neural state moving in the opposite direction along the judgment axis. At the same time, some observations should elicit similar effects on judgment across queries (e.g. observing the queried cause without the effect should have a strongly negative change in judgment regardless of what the queried effect is), and thus the alignment of the left eigenvector relative to some of the inputs should be similar across queries. These two competing demands appear to induce the geometric organization of the line attractors shown in Figure 7 and Figure 6.

### D Limitations

This study represents a first step towards understanding neural mechanisms of causal reasoning. As such, there are a number of important limitations. For example, the variant of the DAG task we have analyzed only considers facilitatory causes, leaving open how neural systems might also reason about inhibitory or mixed (facilitatory and inhibitory) causal effects. Our study also assumes causal interactions follow the disjunctive “noisy-or” parameterization, as opposed to e.g. conjunctive interactions where multiple causes must co-occur to elicit an effect. Further, causal structure learning in cognitive science is believed to also occur through interventions, which we did not explore in this study. Addressing these limitations is an important direction for future work.

### E Causal reasoning in large language models

The rapid development of large language models (LLMs) has generated growing interest in evaluating their causal reasoning abilities [75–80]. For example, Lyu et al. [75] proposed to probe causal judgments by constructing competing natural language causal narratives for pairs of variables *X, Y* (corresponding to *X →Y* versus *Y →X*), and then comparing an LLM’s ability to perform a zero-shot prediction task when prompted under each framing. Early studies of this kind have also motivated the development of systematic benchmarks to evaluate causal reasoning via natural language [81, 82]. However, such approaches differ from our work in several important respects. First, unlike the RNNs studied here, which can be directly analyzed as biologically plausible recurrent dynamical systems, the various mechanisms underlying LLMs (positional embeddings, self-attention, etc) are not straightforwardly aligned with neural computation in the brain. Second, causal reasoning in LLMs is typically elicited through natural language prompts, which likely engage rich semantic and linguistic representations that make it much more difficult to isolate the specific mechanisms that implement causal judgments. By contrast, our proposed DAG task abstracts away from natural language scenarios, enabling a direct analysis of how causal judgments can emerge from recurrent neural dynamics.

### F Biological plausibility of neural mechanisms in RNNs

RNNs have been remarkably successful in generating novel hypotheses for how computations can be implemented via recurrent neural dynamics in the brain. For example, there is a sizeable literature on computation through recurrent dynamics for motor control [83–85], but also increasingly in studies of cognition, e.g. context-dependent decision-making in primates [44] and rats [61], temporal cognition in primates [49], evidence accumulation in primates [86] and spatial navigation [87]. Thus, this modeling approach represents a highly promising methodology for identifying biologically plausible mechanisms underlying cognitive functions.

### G Comparing RNNs with transformers

To compare RNNs against a separate, non-recurrent architecture, we performed an experiment where we trained a set of transformer models (*n* = 5) to perform the DAG task (Figure S3). Interestingly, transformers performed the task accurately (mean accuracy 0.99 across 5 models on the 3-node task), required fewer training examples (just 5000 epochs compared to 25,000 for RNNs), and recapitulated the performance decline when randomizing node *C* during evaluation of query *A →B* (mean accuracy when judging *A →B* in the fork DAG {*C →A, C→ B}*, 0.99; mean accuracy after randomization of *C*, 0.78). These results suggest that certain strategies for causal judgment can be shared across model architectures. We leave a mechanistic analysis of how transformers solve the DAG task for future work.

### H Computational resources

Models were trained on the Axon computer cluster at the Zuckerman Institute at Columbia University using nodes comprised of two Xeon E5-2660 v4 CPUs, eight GTX 1080 Ti GPUs, and 125GB RAM.

### I Code availability

Code used to train the causal RNN models (using PyTorch [88]) is available at: https://github.com/marcustriplett/causalRNN.

## Supplementary figures

**Figure S1:**
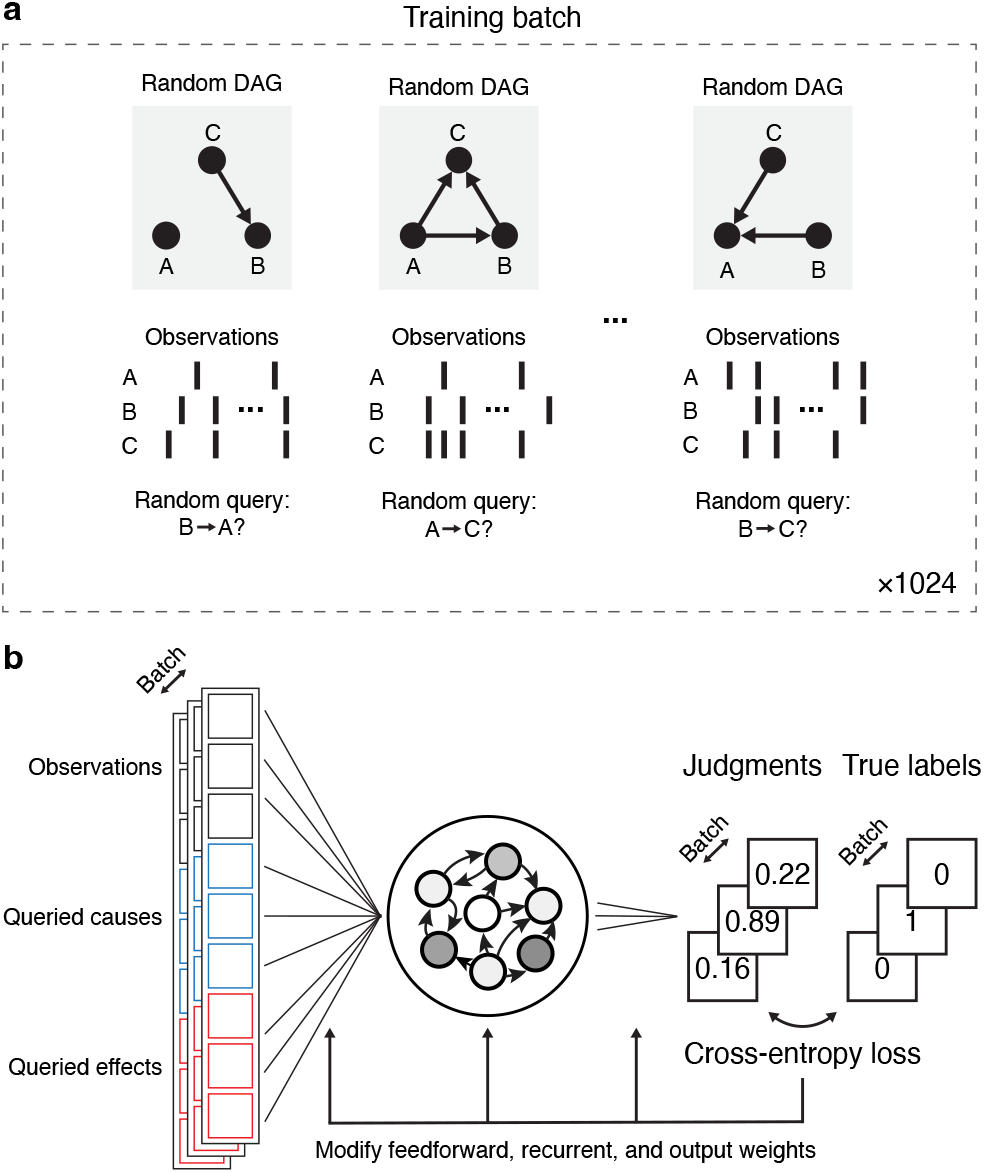
Overview of RNN training paradigm. **a**, A batch consists of 1024 training examples. Each example is made up of a sequence of observations from a randomly sampled DAG, a random query, and the true label (0 or 1 corresponding to whether the query is false or true). **b**, The RNN generates numerical judgments about whether the query is true or false, for all examples in the batch simultaneously, which are compared to the true labels and used to adjust the input, recurrent, and output weights via error backpropagation.

**Figure S2:**
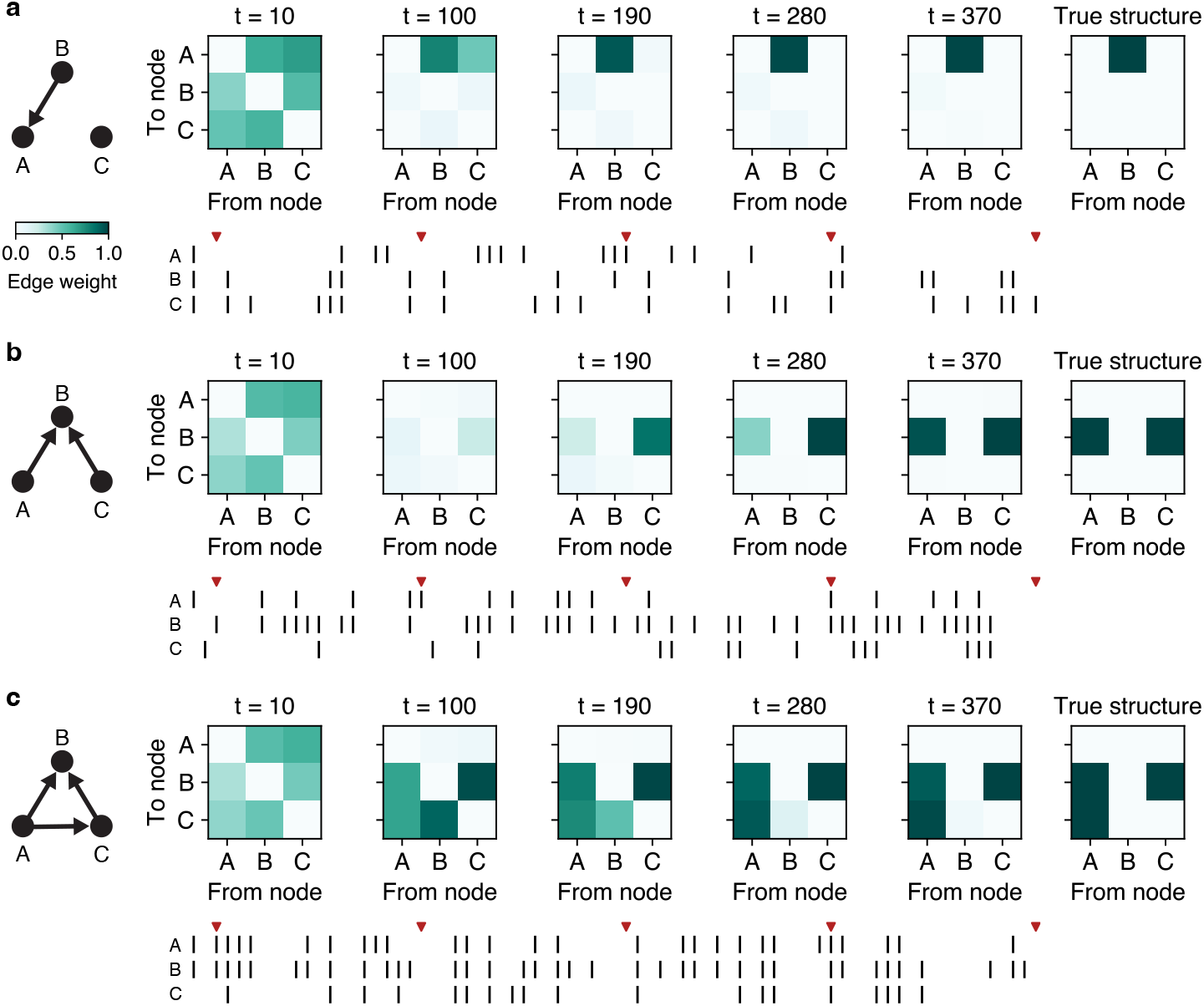
Additional examples of how RNNs’ judgments about entire DAG structures evolve through time, for a singleton DAG (**a**), a collider DAG (**b**), and a mediation DAG (**c**).

**Figure S3:**
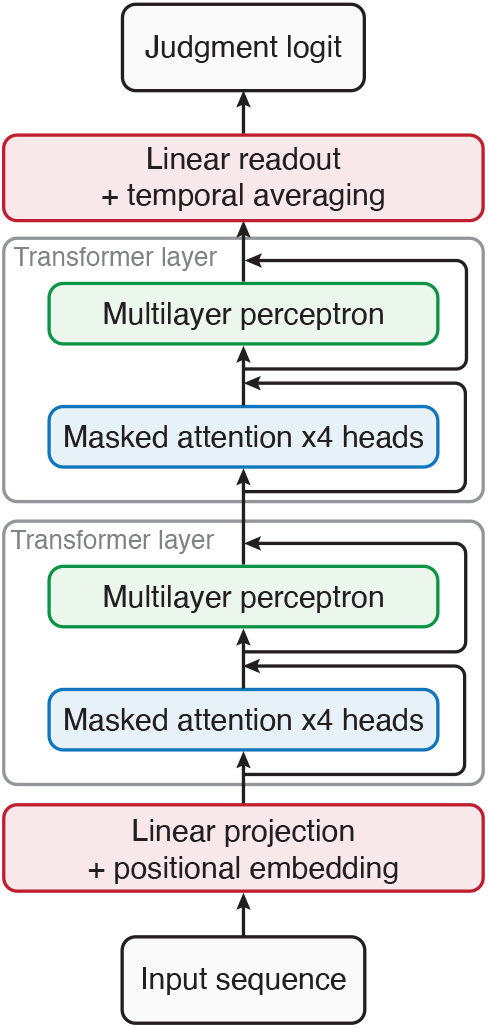
A simple transformer model for causal judgment, comprising a linear projection and positional embedding layer, two transformer layers (4 attention heads per layer, with causal masking), and a linear readout layer.

**Figure S4:**
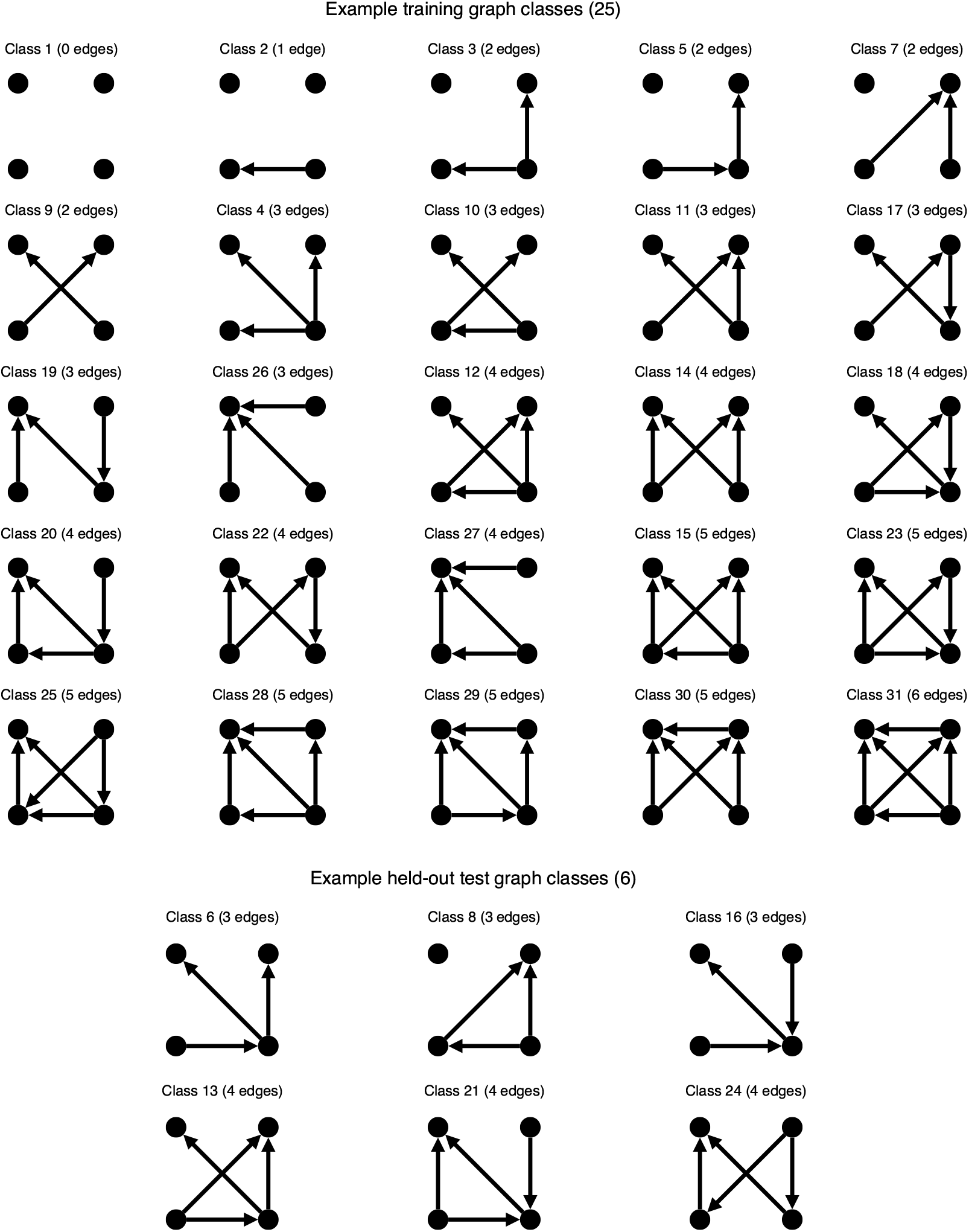
Example partition of the total set of graph classes on 4 nodes into training (top) and test (bottom) sets. Each graph class represents a unique set of causal interactions.

**Figure S5:**
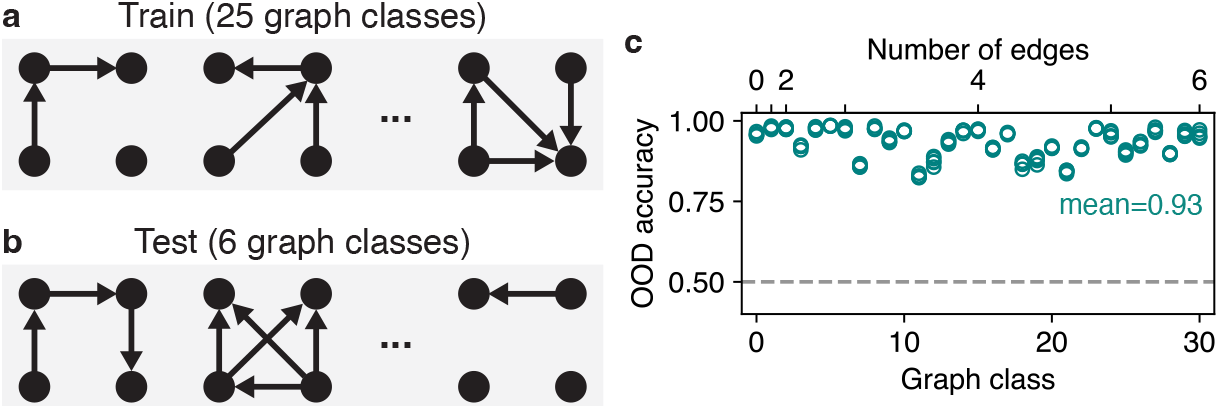
RNNs can generalize to unseen graph classes. **a**, 25 classes of DAGs are used for training. **b**, The 6 classes of DAGs not used for training are held out for testing RNN performance. **c**, RNNs accurately perform the DAG task on the test graph classes (mean accuracy, 0.93). Intriguingly, variability in performance suggests differences in generalization ability for specific training-test splits and graph classes.

**Figure S6:**
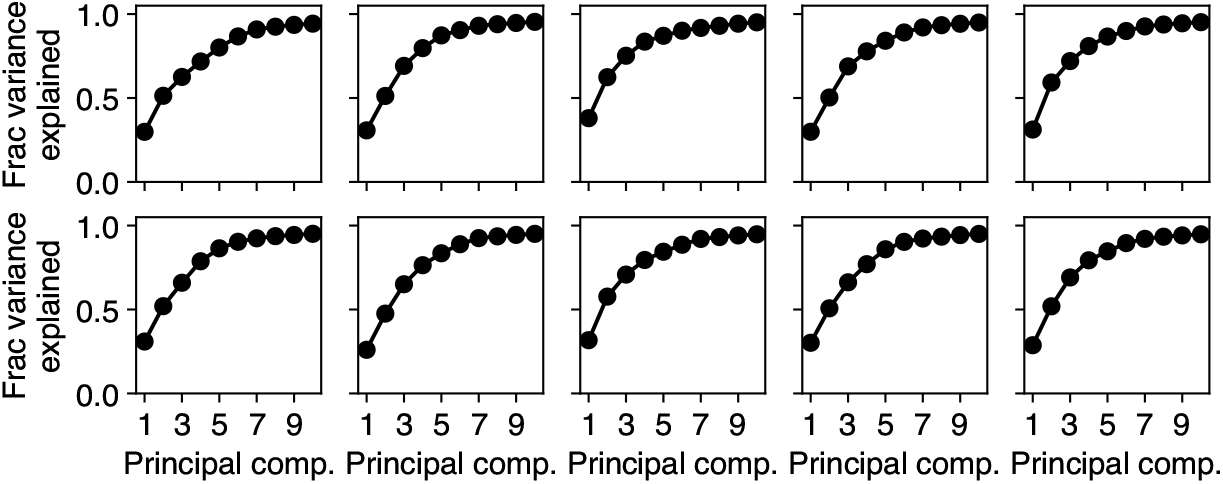
Cumulative variance explained by the first 10 principal components, across 10 randomly initialized RNNs trained on the 3-node DAG task.

**Figure S7:**
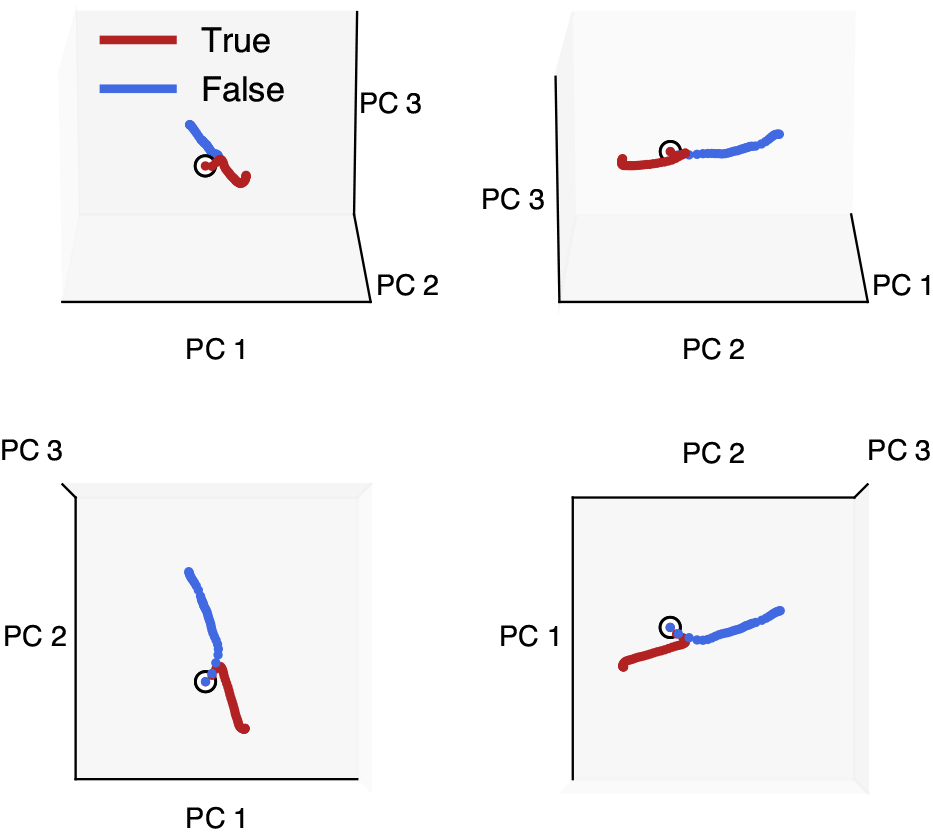
Decision axis (i.e. “judgment axis”), derived from neural activity by averaging trajectories corresponding to DAG edges that exist (“True”, red) vs edges that do not exist (“False”, blue). Averages are taken over 50 samples from each of the 25 possible DAGs on three variables. Cf. Figure 6 for a direct visualization of the decision axis in the top principal components via projection of the RNN readout weights.

**Figure S8:**
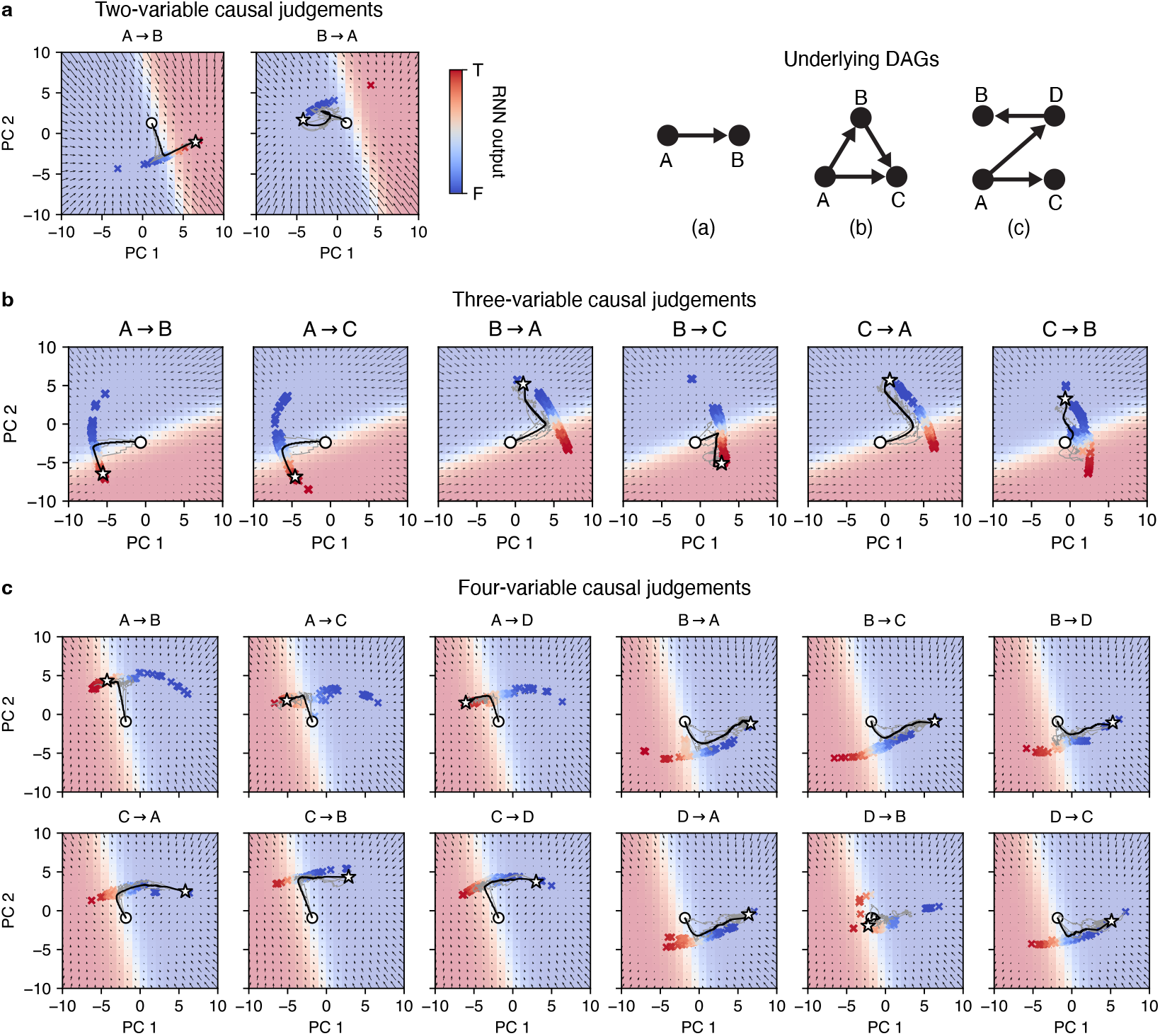
Comparison of fixed point structure from RNNs trained on DAGs having two (**a**), three (**b**), and four (**c**) nodes.

**Figure S9:**
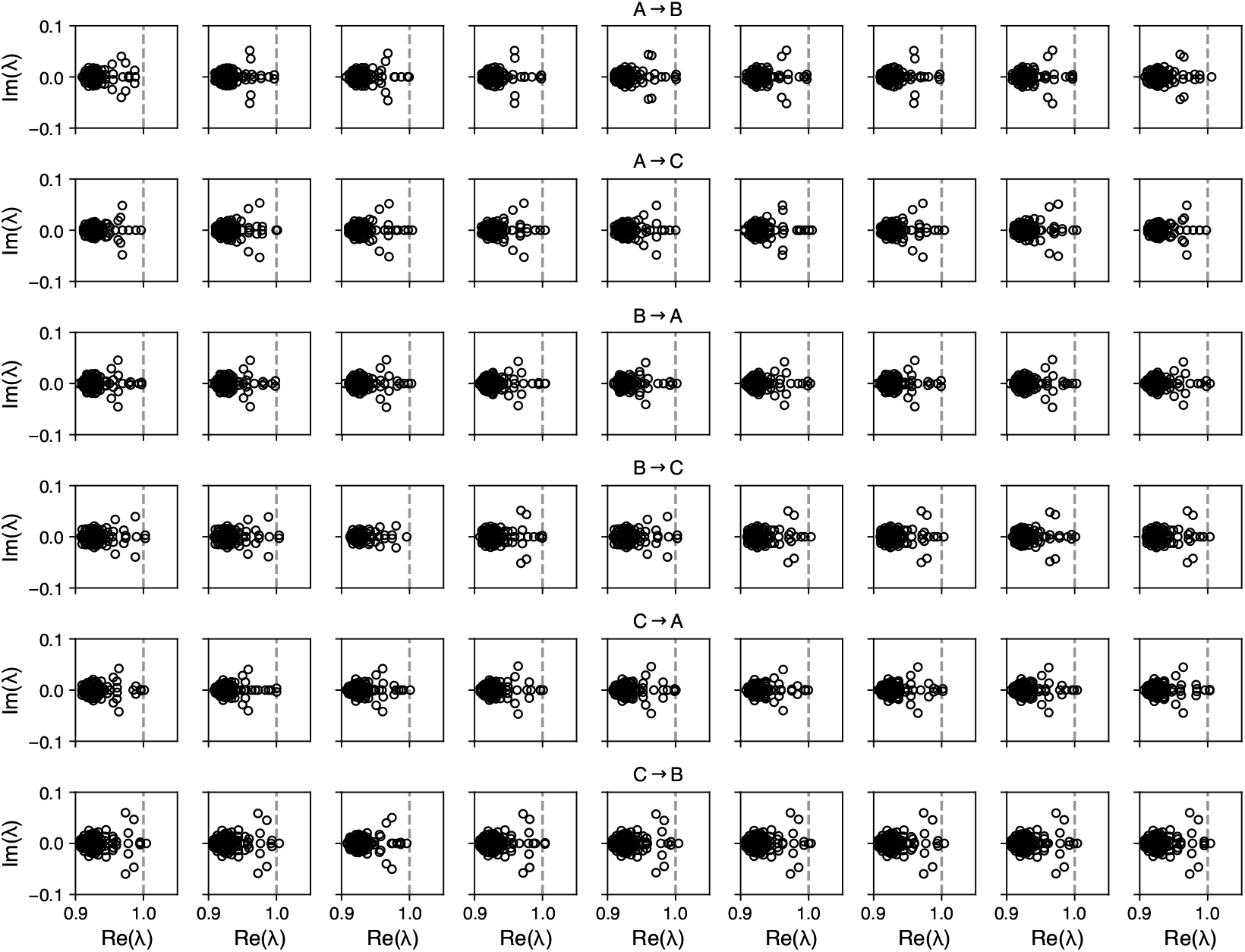
Eigenvalue spectra corresponding to nine randomly chosen fixed points from each of the six line attractors from Figure 5.

**Figure S10:**
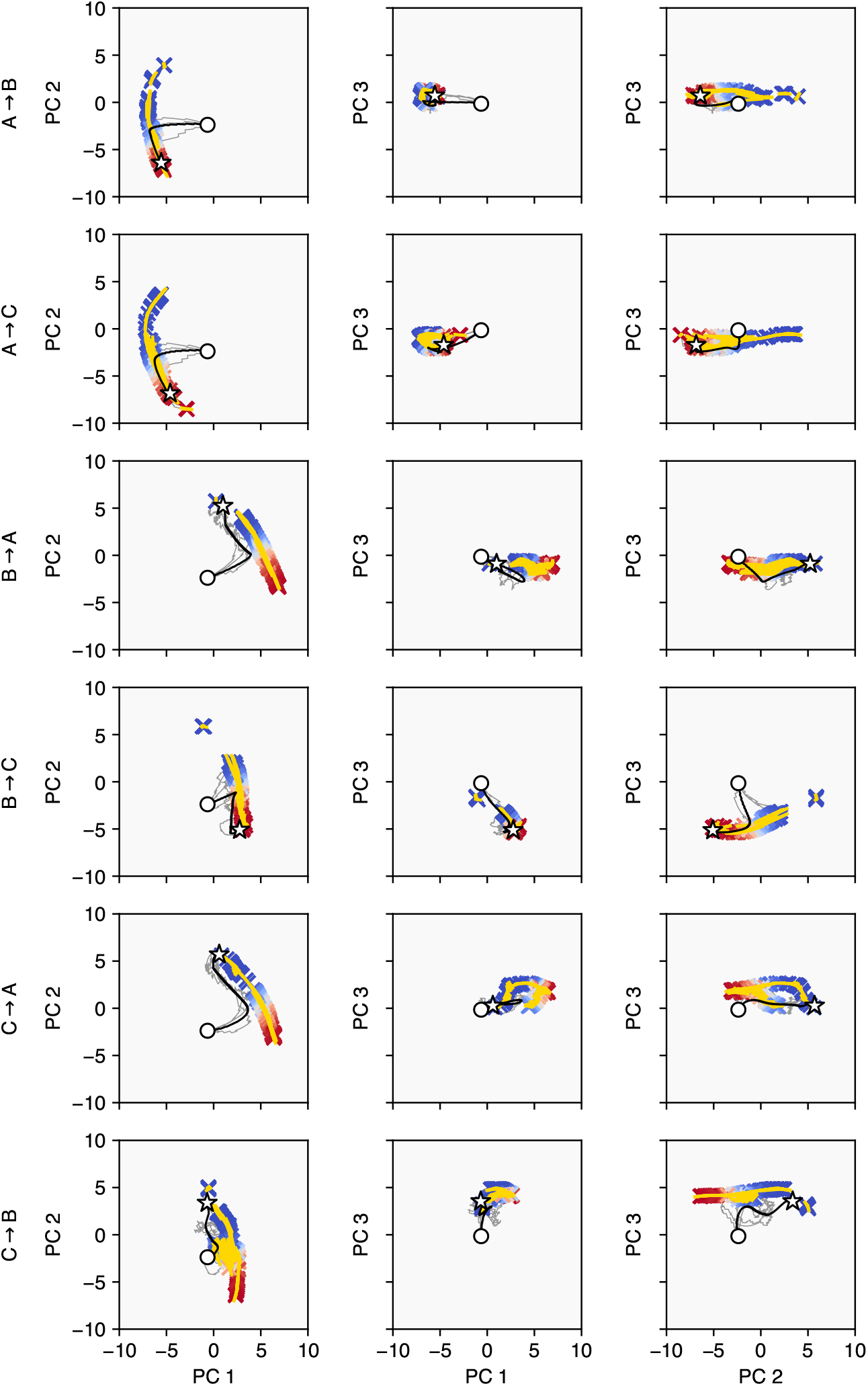
Eigenvectors (yellow lines) associated with the largest real eigenvalues align with the principal direction of the line attractor (red/blue crosses). White circles show neural state at trial start, white stars show neural state at trial end. Black lines show averages over 500 trajectories. Faint gray lines show three example individual trajectories.

**Figure S11:**
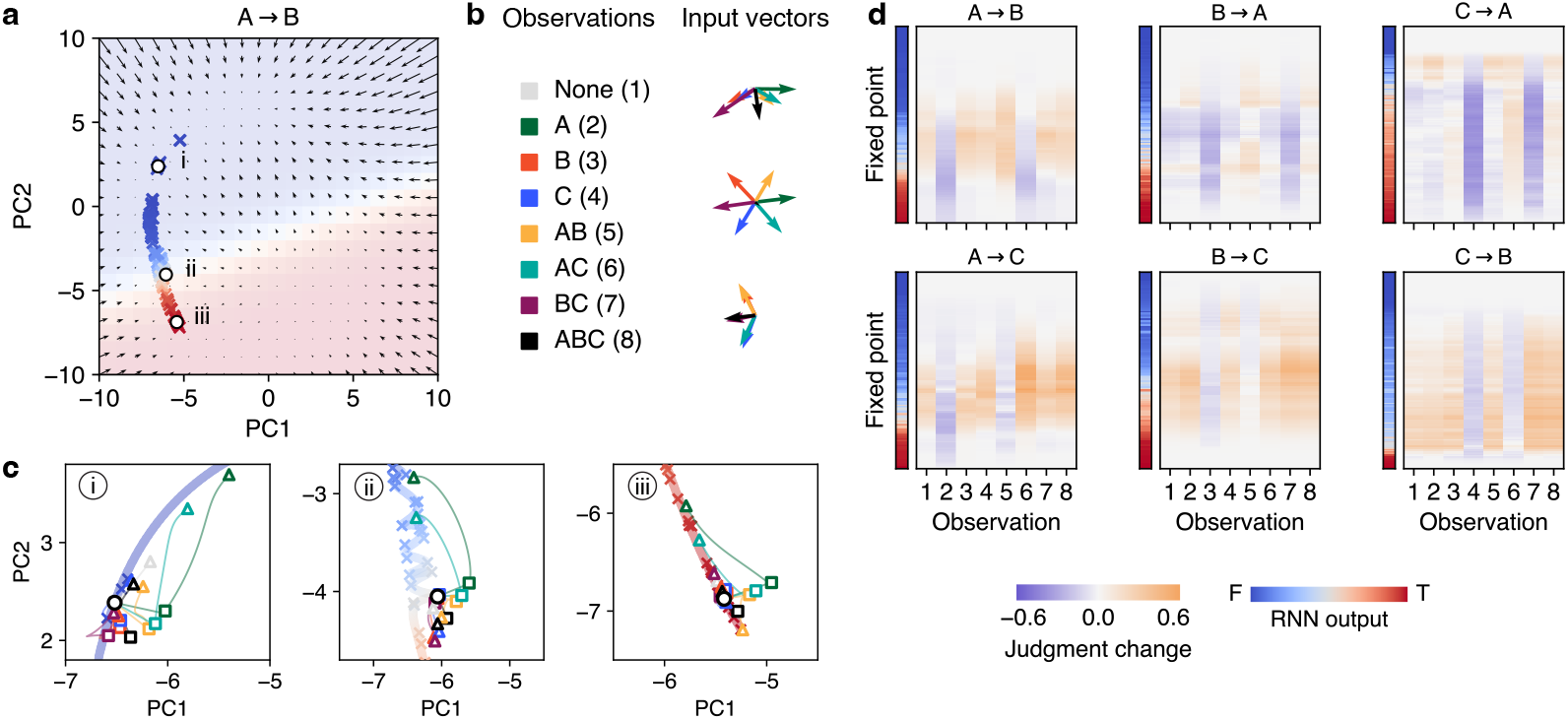
Characterization of how different inputs elicit judgment changes. **a**, Three example fixed points along the *A → B* line attractor. **b**, Left: Colors representing each observation. Right: Input vectors associated with each observation. **c**, Additional examples of the interaction between input vectors and recurrent dynamics, corresponding to the fixed points in (a). **d**, Change in judgment resulting from each observation when the neural state is initialized at a given fixed point. Sufficient time is given for the recurrent dynamics to carry the neural state back to the line attractor. A diversity of state-dependent input effects is revealed through this analysis, that together describe the way in which an RNN makes its judgments. For example, when resolving whether *A* causes *B* in the case of the fork DAG with edges {(*C, A*), (*C, B*)}, observing *AC* almost always moves neural state towards the “false” end of the line attractor. Observing *BC* has a more subtle effect, however, as the observation pushes the neural state towards “true” if the RNN is uncertain, but pushes the neural state back towards “false” if it already believes that *A* does cause *B*. The RNN therefore must rely on instances of *AB* to confirm that *A* causes *B*, whereas instances of *A* and *AC* (the candidate cause without the effect) strongly push the RNN towards “false.”

**Figure S12:**
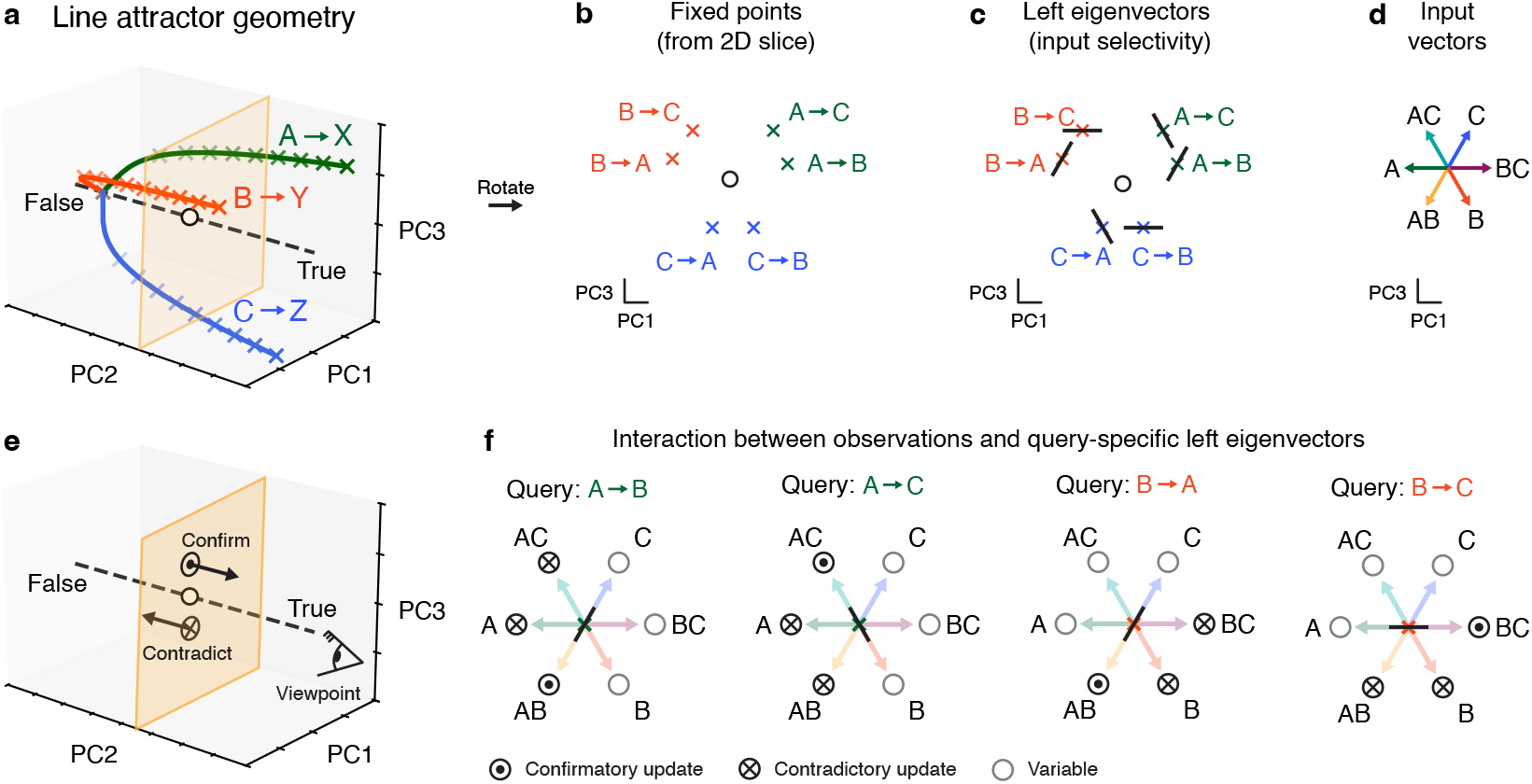
Logical principles of judgment updates in RNNs. **a**, Organization of line attractors. Orange plane shows an example 2D “slice”. **b**, Fixed points associated with the 2D slice in (a). Note that fixed points (and therefore line attractors) are clustered hierarchically: queries with the same cause are closest together, followed by queries with the same effect. **c**, Left eigenvectors (black lines, associated with each fixed point) determine selectivity to observations. **d**, Input vectors associated with each observation. **e**, Schematic of how confirmatory (favoring “true”) vs contradictory (favoring “false”) evidence shifts neural state towards true vs false ends of the line attractor. **f**, Direction of movement along the line attractor (corresponding to network’s change in judgment) is determined by how the input vector aligns with the left eigenvector. As the query changes (e.g. from *A → B* to *A→ C*), the eigenvectors are reconfigured: the orientation of the *left* eigenvector changes to become aligned with observations of *AC*, and anti-aligned with observations of *AB*. This reorientation serves to appropriately push the neural state in confirmatory or contradictory directions (circled dot vs circled cross, respectively). Note that the visual perspective corresponds to that of the cartoon eye in (e).

## Footnotes

1 Notably, simple transformers trained on the same task exhibited similar behaviors (see Appendix G, Figure S3).

2 By convention, we adjust the sign of the right eigenvector so that it aligns with the decision axis.

## Notes

### Competing Interest Statement

The authors have declared no competing interest.

### Summary of Updates

Updated text throughout manuscript; new figure on line attractor geometry added to main text; improvements made to main text figures; additional supplementary figures added.

